# Predicting Alzheimer’s disease progression trajectory and clinical subtypes using machine learning

**DOI:** 10.1101/792432

**Authors:** Vipul K. Satone, Rachneet Kaur, Anant Dadu, Hampton Leonard, Hirotaka Iwaki, Mary Makarious, Lana Sargent, for the Alzheimer’s Disease Neuroimaging Initiative, Ali Daneshmand, Sonja W. Scholz, Andrew B. Singleton, Mike A. Nalls, Roy H. Campbell, Faraz Faghri

**Author notes:** Corresponding author (FF). Data used in preparation of this article were obtained from the Alzheimer’s Disease Neuroimaging Initiative (ADNI) database (adni.loni.usc.edu). As such, the investigators within the ADNI contributed to the design and implementation of ADNI and/or provided data but did not participate in analysis or writing of this report. A complete listing of ADNI investigators can be found at: http://adni.loni.usc.edu/wp-content/uploads/how_to_apply/ADNI_Acknowledgement_List.pdf.

## Abstract

**Background:** Alzheimer’s disease (AD) is a common, age-related, neurodegenerative disease that impairs a person’s ability to perform day-to-day activities. Diagnosing AD is challenging, especially in the early stages. Many patients still go undiagnosed, partly due to the complex heterogeneity in disease progression. This highlights a need for early prediction of the disease course to assist its treatment and tailor therapy options to the disease progression rate. Recent developments in machine learning techniques provide the potential to not only predict disease progression and trajectory of AD but also to classify the disease into different etiological subtypes.

**Methods and findings:** The work shown here clusters participants in distinct and multifaceted progression subgroups of AD and discusses an approach to predict the progression rate from baseline diagnosis. We observed that the myriad of clinically reported symptoms summarized in the proposed AD progression space corresponds directly with memory and cognitive measures, which are routinely used to monitor disease onset and progression. Our analysis demonstrated accurate prediction of disease progression after four years from the first 12 months of post-diagnosis clinical data (Area Under the Curve of 0.96 (95% confidence interval (CI), 0.92-1.0), 0.81 (95% CI, 0.74-0.88) and 0.98 (95% CI, 0.96-1.0) for slow, moderate and fast progression rate patients respectively). Further, we explored the long short-term memory (LSTM) neural networks to predict the trajectory of an individual patient’s progression.

**Conclusion:** The machine learning techniques presented in this study may assist providers in identifying different progression rates and trajectories in the early stages of the disease, hence allowing for more efficient and personalized healthcare deliveries. With additional information about the progression rate of AD at hand, providers may further individualize the treatment plans. The predictive tests discussed in this study not only allow for early AD diagnosis but also facilitate the characterization of distinct AD subtypes relating to trajectories of disease progression. These findings are a crucial step forward for early disease detection. These models can be used to design improved clinical trials for AD research.

## Introduction

Alzheimer’s disease (AD) is a progressive and age-associated neurodegenerative disease affecting a patient’s memory, intellectual skills, and other mental functions. It is the most common form of dementia. Research has shown that AD is a clinically heterogeneous condition, showing marked variations in terms of the symptom constellations and disease progression rates. The clinical signs and symptoms of AD show marked variability in terms of patients’ age, disease span, progression velocity, and types of memory, cognition, and depression-related features. After the age of 65, the prevalence of dementia doubles every five years and is known to increase exponentially after the age of 90 [1]. As dementia affects older people, with a growing life expectancy, it is posing a serious and increasing socioeconomic challenge [2].

With no preventive interventions known, AD progression is a major concern for healthcare providers around the globe. Researchers have shown that AD pathological changes occur 20 years or earlier before the actual disease symptoms manifest [3], [4], [5], [6], [7], [8]. In the absence of any cure or disease-modifying treatment for this disabling disease, current treatment strategies are limited to supportive, symptomatic care [9], [10]. Delay in the diagnosis of AD is often due to the disease complexity, with limited early diagnostic metrics available for healthcare providers [11]. A major challenge for AD prediction is the presence of inherent phenotypic diversity in the AD population, limiting diagnosis, prognosis, and counseling of affected patients regarding their individual risks and expected progression rate. This problem becomes particularly burdensome as we move increasingly toward early-stage clinical trials when therapeutic interventions are likely to be most effective. The ability to predict and account for even a proportion of the disease course can significantly reduce the cost of clinical trials and increase the ability of such trials to detect treatment effects. Therefore, predicting disease progression trajectories at an early stage is crucial for the design of clinical trials and the development of disease-modifying treatment strategies.

For the treatments to be most effective, the AD therapy regimen must likely begin before notable downstream damage occurs [12]. Simply put, early AD detection is a likely scenario to yield the greatest therapeutic gains. Patients diagnosed with amnestic mild cognitive impairment (MCI) at study baseline are at a higher risk for progression to dementia, but not all patients end up developing AD [13]. Research has been done to detect AD in patients with MCI or predict the early stage of AD using cerebrospinal fluid (CSF) biomarkers [14], [15], while others [16] have used psychometric and imaging data for predicting the progression of dementia in patients with amnestic MCI. In an implementation of a multiclass classifier using clinical and magnetic resonance (MR) brain images to classify controls, MCI and AD patients, an accuracy of 79.8% was achieved [17]. Less research has been done on using only clinical data to predict the AD progression rate. Faghri *et al*. [18] used machine learning to classify Parkinson’s disease (PD) patients into three different sub-categories with highly predictable progression rates. They explored variations in onset and progression velocity and observed clusters of the motor, cognitive and sleep disturbance related features using only clinical data. Here, we extend this approach by applying it to the clinical features of the Alzheimer’s Disease Neuroimaging Initiative (ADNI) dataset. This study is in continuation of the investigation regarding early AD onset and progression started in [19].

### Goals and Contributions

This study is designed to describe and predict the clinical progression of AD at an early stage. The first stage of this effort requires creating a multi-dimensional space that captures both the features of the disease and the progression rate of these features (i.e., velocity). Rather than creating a space based on *a priori* concepts of differential symptoms, we used data dimensionality reduction methods on the complex clinical features observed at 24 and 48 months after initial diagnosis to create a meaningful spatial representation of each patient’s status at this time point. After creating this space, we used unsupervised clustering to determine whether there were clear subtypes of disease within this space. This effort delineated three distinct clinical subtypes corresponding to three groups of patients progressing at varying velocities (i.e., *slow, moderate*, and *fast* progressors). Following the successful creation of disease subtypes within a progression space, we created a baseline predictor that accurately predicted an individual patient’s clinical group membership four years later. We have also developed a machine learning model to predict an individual’s progression trajectory. This highlights the utility of machine learning as ancillary diagnostic tools to identify disease subtypes and to project individualized progression rates based on model predictions.

This study was designed to cluster AD patients into distinct progression groups and to predict the progression trajectory at an early baseline period. Dimensionality reduction via non-negative matrix factorization (NMF) was used to define an *ADNI progression space* for the AD summarizing myriad clinical measures across multiple time points. By applying unsupervised machine learning, namely, the Gaussian mixture model (GMM) on the extensive clinical observations available in the ADNI dataset, we algorithmically parsed the progression space for AD into three clinical subtypes, defined as *low, moderate (medium)* and *high* disease rate progressors. Our analysis found that clinically related measures corresponding to memory and cognition make up the AD progression space. Clinical data collected at baseline (study entry), after 6 and 12 months, was used to predict memory and sleep decline after 24 and 48 months from baseline. We validated our models through five-fold cross-validation to obtain a robust prediction of memberships into these progression subtypes. Along with traditional machine learning methods, the long short-term memory (LSTM) neural networks were also used to predict disease progression rates (control, low, moderate, and high) after 24 and 48 months from baseline. The described methodologies pave a way forward towards the development of personalized clinical care and counseling for patients, hopefully reducing AD therapy costs in the future.

Further, we examined the reversion instances of AD captured in the constructed progression space, the correlation of Apolipoprotein Eε4 (APOEε4) compound genotype with cognitive performance and interactions between certain selective features associated with AD and the constructed progression space later in the discussion section of the paper. These observations provide a promising understanding of AD characteristics useful for studying g novel disease modification therapies. We believe that the advancement of the discussed prediction models has the potential to impact clinical decision making and improve healthcare resource allocation in AD significantly.

## Materials and Methods

The data analysis pipeline for this work was performed in Python 3.6 with the support of several open-source libraries (TensorFlow, scikit-learn, pandas, seaborn, etc.). To facilitate replication and expansion of this study, the Python code (including the entire data preprocessing and machine learning analysis) was made publicly available under GPLv3 as part of the supplementary information at https://github.com/vipul105/Alzheimers_Disease_Progression.

### Study design and participants

Data used in the preparation of this article were obtained from the ADNI database (adni.loni.usc.edu). The ADNI was launched in 2003 as a public-private partnership, led by Principal Investigator Michael W. Weiner, MD. The primary goal of ADNI has been to test whether serial magnetic resonance imaging (MRI), positron emission tomography (PET), other biological markers, and clinical and neuropsychological assessments can be combined to measure the progression of MCI and early AD. The ADNI dataset involves participants from over 50 sites across North America and Canada. All participants and their study partners provided their consent, accepting their engagement for the data collection. The study protocols for ADNI were approved by the Institutional Review Board. The ADNI study was carried out in phases, namely, ADNI 1 beginning in 2004, followed by ADNI GO in 2009, ADNI 2 in 2011, and ADNI 3 in 2016. These editions had different participants and data collection procedures, accounting for advancement in technologies. For more up-to-date information, see www.adni-info.org. ADNI 2 study is the main cohort in our study. ADNI 2 is the longest study, with the highest data heterogeneity and availability.

The key eligibility criteria for the ADNI participants are highlighted in the supplementary section S1, and further details on the protocol can be found on the ADNI’s procedure manual [20]. All participants went through comprehensive functional, cognitive, and clinical assessments and provided a blood sample for APOE genotyping at their baseline visit (study entry). These assessments and their status (control, MCI, and AD) were then updated longitudinally at 6, 12, 18, 24, 36 and 48 months. In our analysis, predictions were made for each participant’s AD stage after 24 and 48 months using up to 12 months of clinical data. The study consisted of 247 observations (with 123 [49.79%] females), the average age for all participants was 71.55 ± 6.79 years and 94.73% of them are of European ancestry) for prediction at the 48^th^ month and 453 observations (with 208 [45.92%] females), the average age for all participants was 72.32 ± 7.13 years and 93.59% of them are of European ancestry) for prediction at the 24^th^ month. For observations corresponding to the 24^th^ month, mean age was 72.84 ± 6.09, 71.61 ± 7.47 and 72.92 ± 8.11 for controls, MCI and dementia patients respectively and for observations corresponding to the 48^th^ month, the mean age was 72.17 ± 6.67, 71.36 ± 6.67 and 70.34 ± 7.42 for controls, MCI and dementia patients, respectively.

The total scores and subscores from the following commonly collected cognitive, functional, and longitudinal clinical data elements were used in the proposed work:

1. Montreal cognitive assessment (MoCA; version 8.1) [21]
2. Clinical dementia rating [22]
3. Neuropsychiatric inventory questionnaire [23]
4. Neuropsychological battery [24]
5. Mini-mental state exam (MMSE) [25]
6. Geriatric depression scale [26]
7. Everyday cognition - study partner [27]
8. Everyday cognition - participant [28]
9. Functional assessment questionnaire (FAQ) [29]
10. Alzheimer’s Disease Assessment Scale-Cognitive (ADAS-Cog-11/ADAS-Cog-13) [30,31]

We considered a total of 147 clinical variables (features) from the above-mentioned assessments for our analysis. An elaborated list of features used in each test with their definition is given in the supplementary materials (S2 Table).

### Procedures and statistical analysis

Only the observations which had data recorded for all the considered tests were taken into account. To construct the AD progression space, we used readings taken at baseline and on visits after 6 and 12 months from the baseline.

### ADNI progression space and the prediction model

We leveraged the temporal information present in the data to manage missing data recordings. Missing values were imputed using linear interpolation based on the past visit readings for the feature, therefore avoiding any influence of other observations during data imputations. After the imputation, around 7% of the data was reduced. A descriptive plot with the number of observations available for each feature before and after data imputation is given in the supplementary section S4. One hot encoding was used for categorical variables whenever required. Scaling the continuous features to a comparable range is necessary to avoid the influence of certain features over others. Min-max normalization was used to retain the progressions since the ADNI dataset in consideration is multimodal. Furthermore, min-max normalization did not affect categorical features. Fig 1 shows our detailed workflow pipeline followed during the analysis. To reduce the dimensionality of the dataset, NMF [32] (with a rank of 2) was used on 582 observations with available data for baseline, visits after 6 and 12 months. We used NMF to deconstruct data into two matrices, namely progression vectors and the progression indicators which correspond to the latent vectors. Progression vectors were used to construct the 2-dimensional (2D) ADNI progression space. This 2D space was then used to predict a participant’s disease progression stage after 24 and 48 months from baseline. Progression indicators map the features in the original dataset to the progression space, via which we identified memory and cognitive decline as the two dimensions of the modeled AD progression space. The relative position along the x- and the y-axis represent worsening sleep or memory disorder.

**Fig 1.**
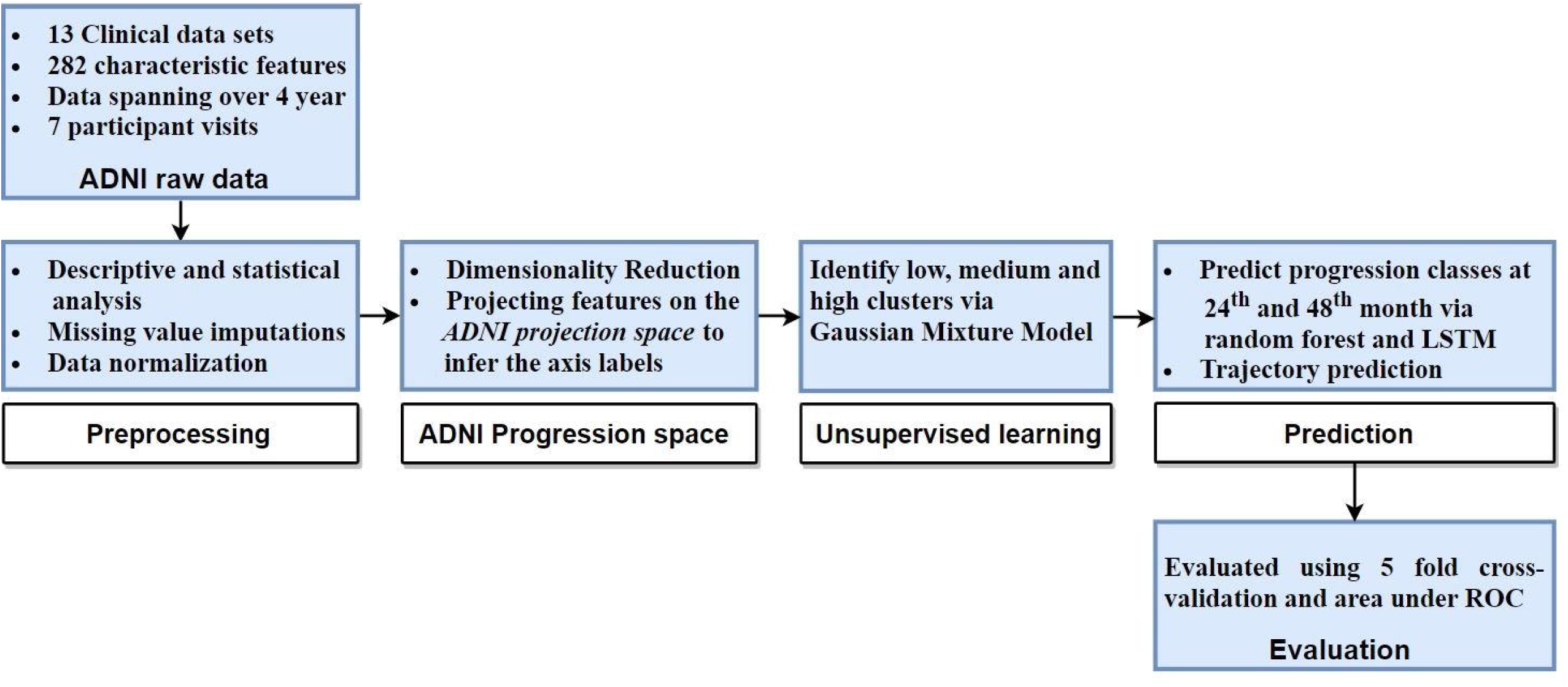
The workflow of analysis and model development.

Next, unsupervised clustering via GMM [33] was used to define the hidden subtypes within the MCI and dementia patients. GMM is an expectation-maximization algorithm that maximizes the likelihood of observing the data, given the underlying parameters of the distribution. Bayesian information criterion (BIC) [34] was used to select the optimum number of underlying clusters for the GMM. BIC is a maximum likelihood estimate which tries to select the best model among the given set of candidates. In all the scenarios, three optimum number of clusters, defined as *low, moderate (medium)* and *high* progression rates, were attained. After obtaining the AD progression space and classifying MCI and dementia patients into different progression groups, the performance of various supervised learning classifiers (namely ensemble random forest, linear discriminant analysis, Naive Bayes, adaptive boosting, nearest neighbors, logistic regression and decision trees) were compared to predict a participant’s progression stage after 24 and 48 months from baseline using readings up to 12 months. Two models were built a) Model 1: predicts progression at 24^th^ month after baseline by using baseline and first-year factors b) Model 2: predicts progression at 48^th^ month after baseline by using baseline and first-year factors. Recurrent neural network (RNN) architecture with LSTM was also used to predict the progression rates (slow, moderate and fast) after 24 and 48 months from the baseline. The LSTM architecture had a single LSTM bidirectional layer connected to a fully connected layer. Cross entropy loss function was used at the output layer since it combines both logs of softmax and negative log-likelihood loss functions. Optimal parameters for the models were found to be a single hidden layer with 128 hidden units with a learning rate of 0.001 and a dropout probability of 0.2. Since our dataset size was limited in terms of the number of observations, 5 fold cross-validation (CV) was used to evaluate the models. Among all the explored algorithms, a random forest classifier [35] gave the best 5 fold CV accuracy. Hence, parameters for the random forest algorithm were fine-tuned using grid search (4800 iterations) and 5 fold CV.

### Model evaluation

Sensitivity and specificity are measures of the proportion of positives that are correctly identified and negatives that are correctly identified, respectively. The plot of sensitivity on the y-axis and 1-specificity on the x-axis is called the area under the receiver operating characteristic (AUC of ROC) curve with a greater value representing a better clustering model. AUC of ROC was used to evaluate the clustering algorithms. Since this is a multiclass problem, one versus all approach was used to calculate the AUC for each class. Next, five-fold cross-validation was used to judge the performance of the proposed prediction models. The model was repeatedly trained on four parts and accuracy for prediction was calculated on the fifth part with a random selection of partitions each time.

### Trajectory prediction

NMF was used to project the observations in progression space. LSTM was used to predict the position of patients in the progression space (trajectory prediction) at the 24^th^ and 48^th^ month using data collected at baseline and after 6 and 12 months from baseline. For this study, five separate projections were made using data until each visit and projections at the 24^th^ and 48^th^ month were predicted. A bidirectional LSTM with 2 layers consisting of 32 hidden units was trained for 50 epochs with a learning rate of 0.001 and a batch size of 10 for the same. Since the projection was done in the 2D axis, the mean squared Euclidean distance was used to assess the performance of this model. Only the features which were present for all the first three visits (baseline, six-month, and 12-month visits) were considered for this study. We used 59 intersecting features in trajectory prediction as opposed to 81 features used in the supervised classification of progression subtypes.

In the subsequent analysis, we studied the share of different frequencies of APOEε4 variants for each progression subtype since APOEε4 genotype is known to be closely related to AD risk [36]. Further, we discuss the reversion from AD to MCI and MCI to control stage captured in the proposed progression space and correlation of a participant’s AD progression stage with their age, educational status, APOEε4 gene, and other selective critical features.

## Results

We have two progression indicator vectors in the reduced 2D ADNI progression space. The features observed in the real data were correlated to the two axes of the progression space using the magnitude of coefficients observed in the progression indicator vectors. A higher magnitude corresponding to the first progression indicator vector will correlate the feature to the first axis and similarly for the second axis.

### ADNI progression space

The observed axis labels for the features using 2D NMF and GMM are presented in the supplementary materials (S3 Table). Progression indicator, i.e., coefficient matrix obtained from the NMF, was used to find out the hidden features that each of the two axes of the reduced space represents (as depicted in section S5 of the supplementary information). Progression indicator vectors represent latent features of the reduced progression space. Progression indicator coefficients for each feature are plotted in Fig 2, and they are separated by drawing a line with slope 1. Features that occur below this separating line were associated with cognitive decline (x-axis) in the AD projection space and features that lie above the line were associated with memory decline (y-axis) in the AD progression space. Features close to the separating line were not associated with any axis.

**Fig 2.**
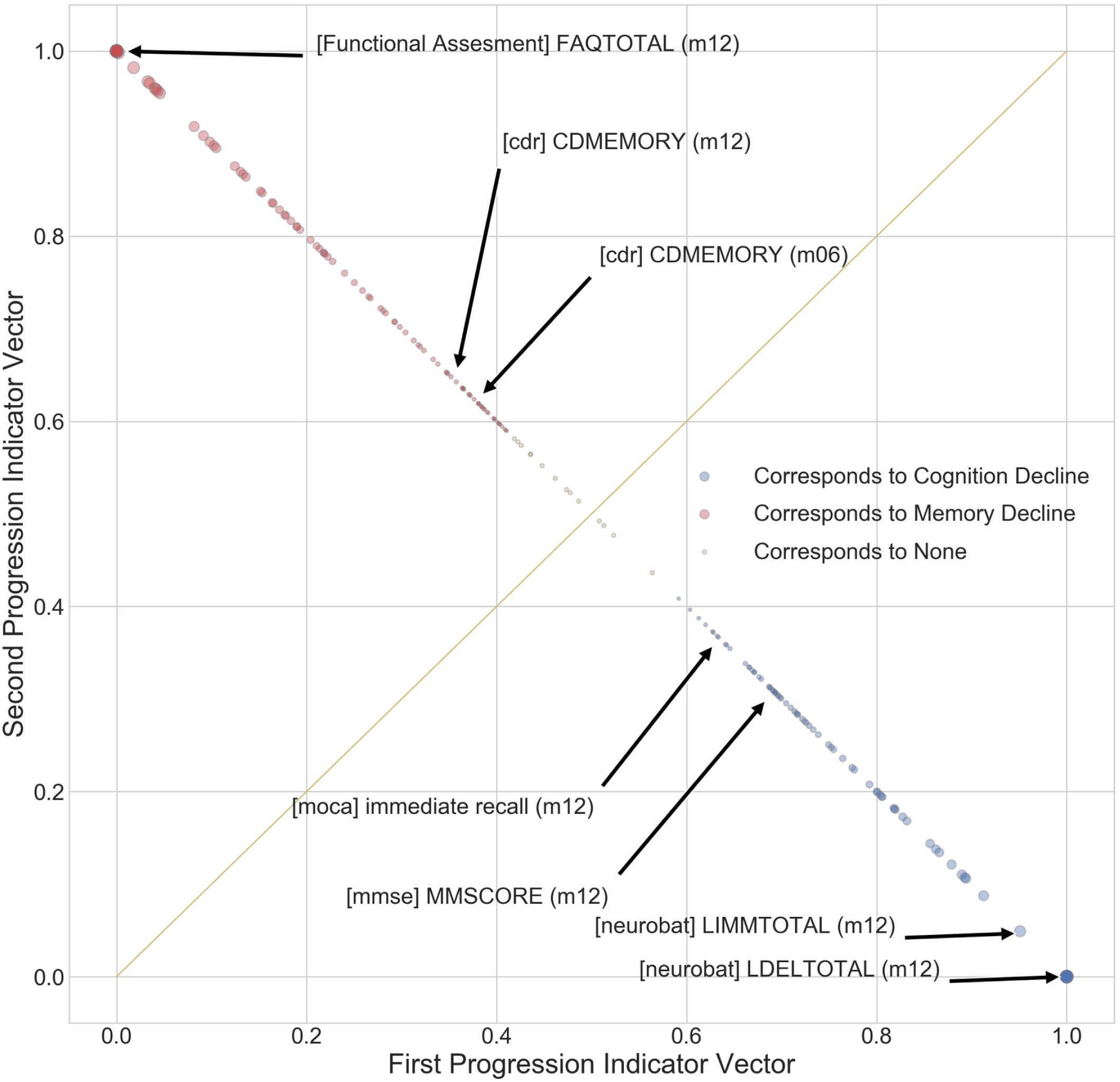
The plot of features in two dimensions using progression indicator vectors. Features in red correspond to memory decline and features in blue correspond to cognitive decline. Yellow line with a slope of 1, which separates the features into two categories, is drawn for reference.

This transformed data was used to project the participant’s disease progression stage at the 24^th^ (Fig 3) and the 48^th^ month (Fig 4).

**Fig 3.**
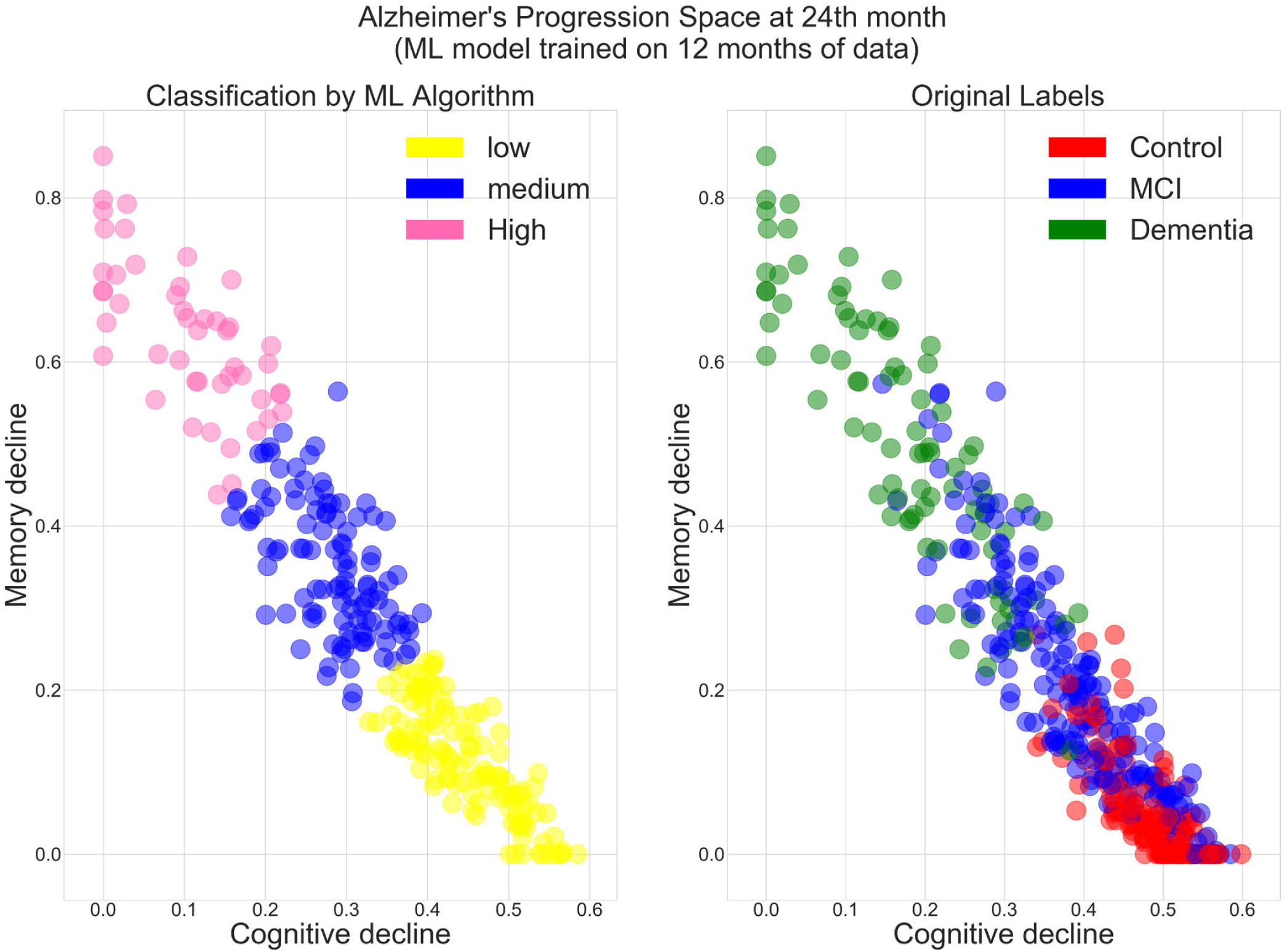
Comparison of 24th month machine learning based prediction and original labels. A total of 582 cases are projected in the AD progression space at the 24^th^ month. **Left:** Machine learning based classification. Low, moderate and high progression rates are represented in yellow, blue and pink, respectively. **Right:** Colored with original labels. Controls, MCI, and dementia patients are represented in red, blue and green respectively.

**Fig 4.**
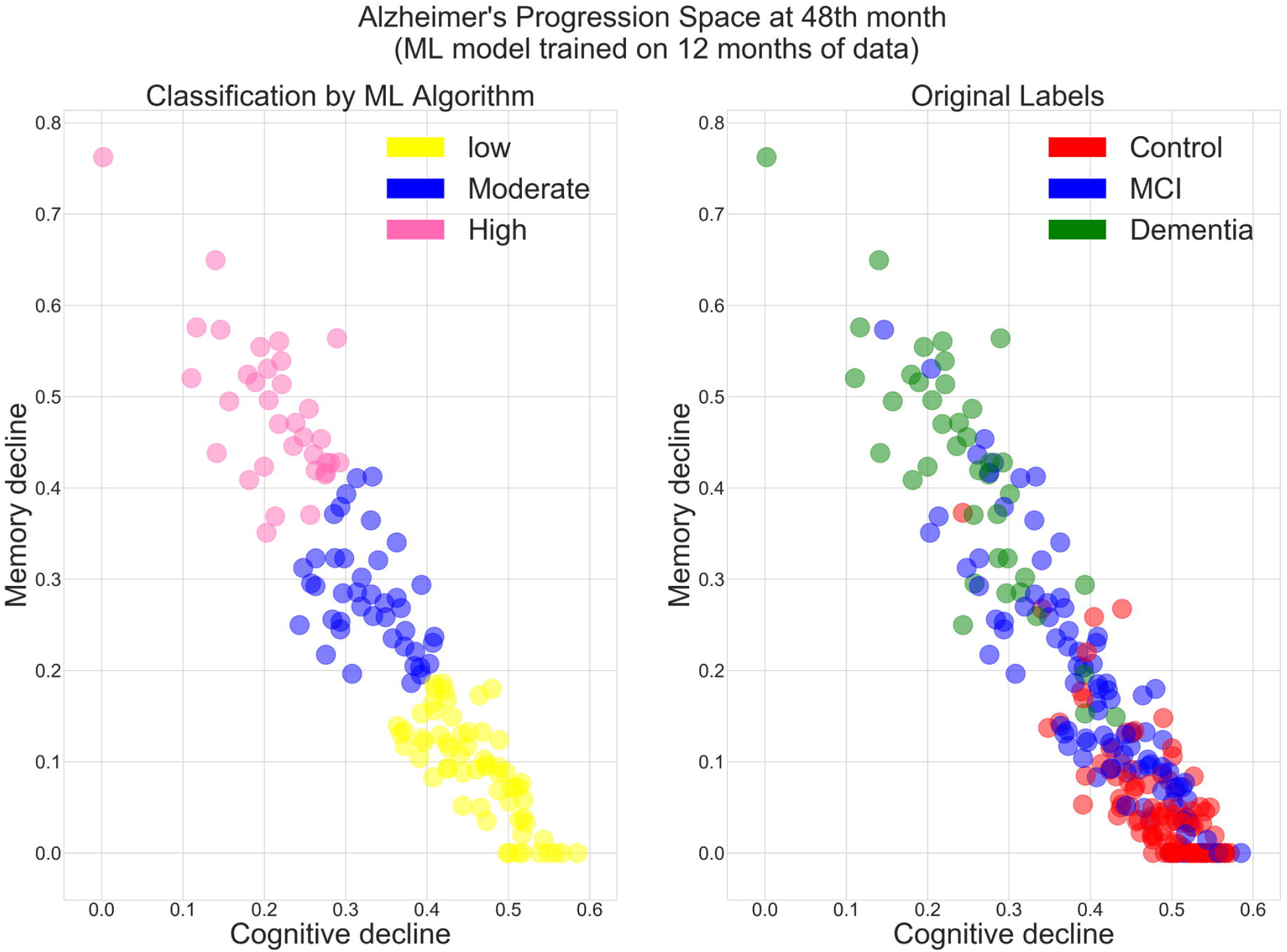
Comparison of 48th month machine learning based prediction and original labels. A total of 253 cases are projected in the AD progression space at the 48^th^ month.**Left:** Machine learning based classification. Low, moderate and high progression rates are represented in yellow, blue and pink, respectively. **Right:** Colored with original labels. Controls, MCI, and dementia patients are represented in red, blue, and green respectively.

Further, three different zones, namely low, moderate, and high progression rates, were identified in the MCI and dementia patients at 24^th^ and 48^th^ month, as depicted in Fig 5.

**Fig 5.**
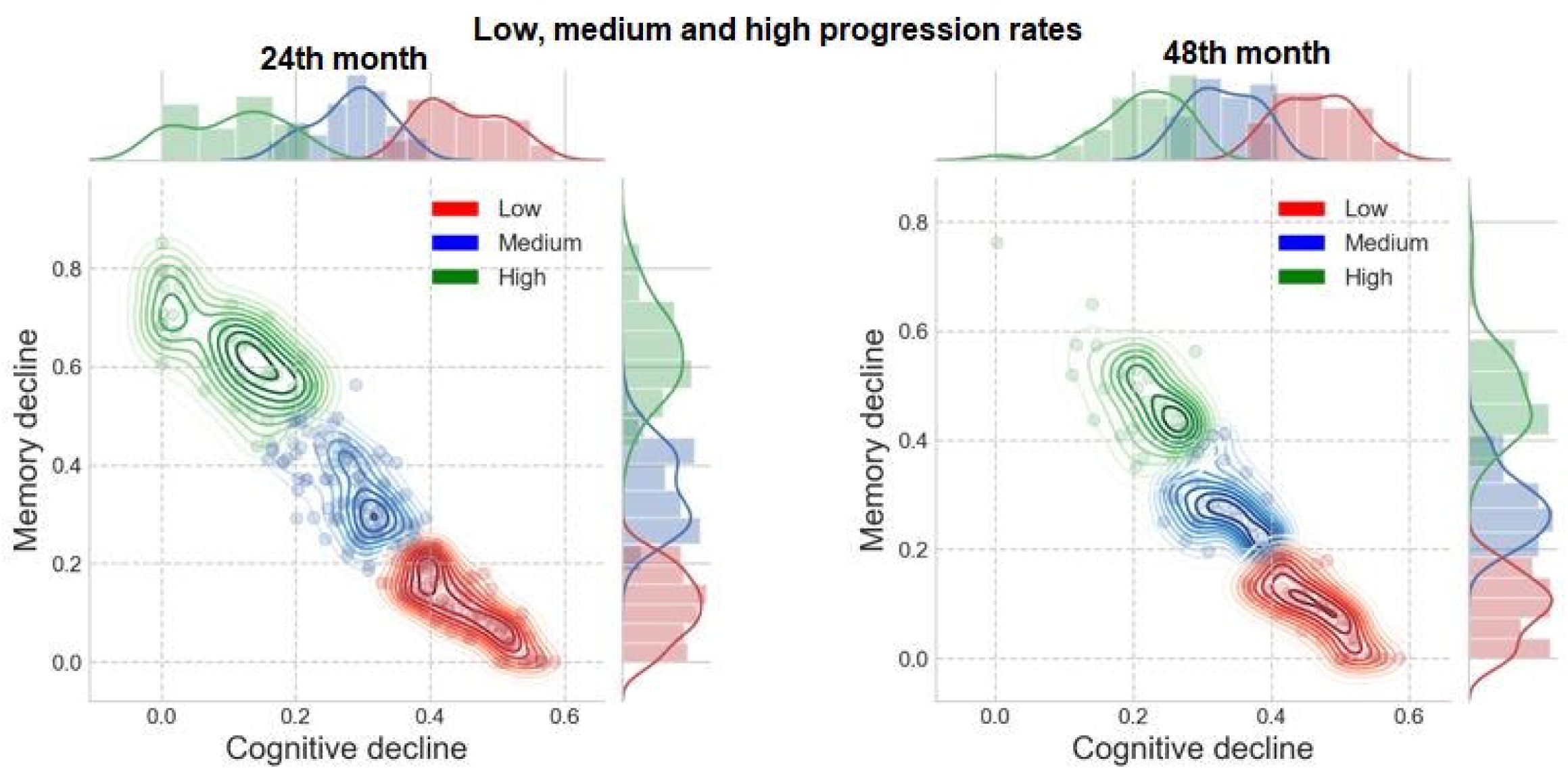
Three different progression rates are identified in MCI and dementia patients. **Left:** at the 24th month. The low, moderate and high progression rate zones are represented in red, blue and green respectively. **Right:** at the 48th month. The low, moderate and high progression rate zones are represented in red, blue, and green respectively.

The progression rate of participants after the 24^th^ and 48^th^ months from the baseline were predicted using the random forest classifier. It gave the best 5 fold cross-validation accuracy and area under ROC curve results for all the cases. A comparison of different machine learning algorithms for the prediction of progression after 24 and 48 months from the baseline is given in the supplementary sections S6 and S7, respectively. For the 24^th^ month using baseline and 12 months of observations, AUC of 0.98 (95% CI, 0.97-0.99), 0.92 (95% CI, 0.89-0.95), and 0.99 (95% CI, 0.98-1.0) for slow, moderate and fast progression rates were observed respectively. Prediction of progression at 48^th^ month using baseline and 12 months of observations yields AUC of 0.96 (95% CI, 0.92-1.0), 0.81 (95% CI, 0.74-0.88) and 0.98 (95% CI, 0.96-1.0) for slow, moderate and fast progression rates respectively. In our implementation, the accuracy for the prediction of progression at the 24^th^ and 48^th^ month is 92.45% and 85.48% respectively (Table 1). Fig 6 depicts the ROC curves for 3 separate classes (slow, moderate and fast progression rates) for the 24^th^ and 48^th^ month.

**Table 1.** Results for model 1 and model 2.

| Progression Month | Accuracy<br>(95% CI) | AUC<br>(95% CI) |  |  |
| --- | --- | --- | --- | --- |
|  |  | Slow | Moderate | Fast |
| **Prediction at M24 using Baseline data** | $0.8113 \pm 0.0$ | $0.94 \pm 0.03$ | $0.84 \pm 0.06$ | $0.94 \pm 0.04$ |
| **Prediction at M48 using Baseline data** | $0.83064 \pm 0.01619$ | $0.94 \pm 0.03$ | $0.75 \pm 0.07$ | $0.95 \pm 0.03$ |
| **Prediction at M24 using one year of data** | $0.9245 \pm 0.0069$ | $0.98 \pm 0.01$ | $0.92 \pm 0.03$ | $0.99 \pm 0.01$ |
| **Prediction at M48 using one year of data** | $0.8548 \pm 0.0$ | $0.96 \pm 0.04$ | $0.81 \pm 0.07$ | $0.98 \pm 0.02$ |

**Fig 6.**
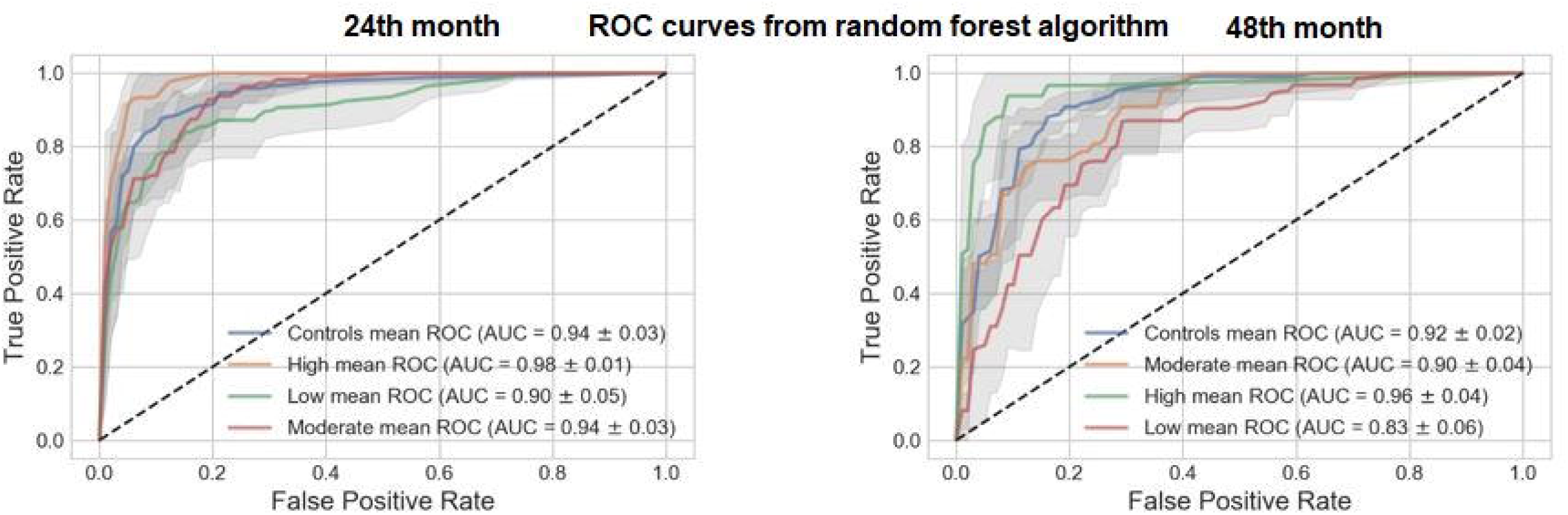
ROC of AD patient’s progression rate after the 24th and 48th months based on the baseline data. Including the area under the ROC for 3 AD progression subtypes (slow, moderate and fast progression rates). **Left:** The predictions for the disease stage at the 24^th^ month were made using a random forest algorithm. **Right:** The predictions for the disease stage at the 48^th^ month were made using random forest algorithm

LSTM model was also used for the prediction of AD subtypes with low, medium and high progression rates as well as controls. The accuracy of prediction of projection rates using LSTM is 82.58 +/- 1.58% and 87.40 +/- 1.38% for the 48^th^ and 24^th^ month respectively. The performance of the neural network did not match other more traditional methods because of small sequence length, a smaller number of features. and a limited number of participants in the dataset.

### Trajectory prediction using LSTM networks

Since no therapies in AD modification are known yet, predicting the trajectory of AD progression at the early disease stage may offer a practical tool for refined clinical trials testing of novel disease-modifying strategies. For example, Fig 7 represents the predicted and projected values in the AD progression space for one of the participants.

**Fig 7.**
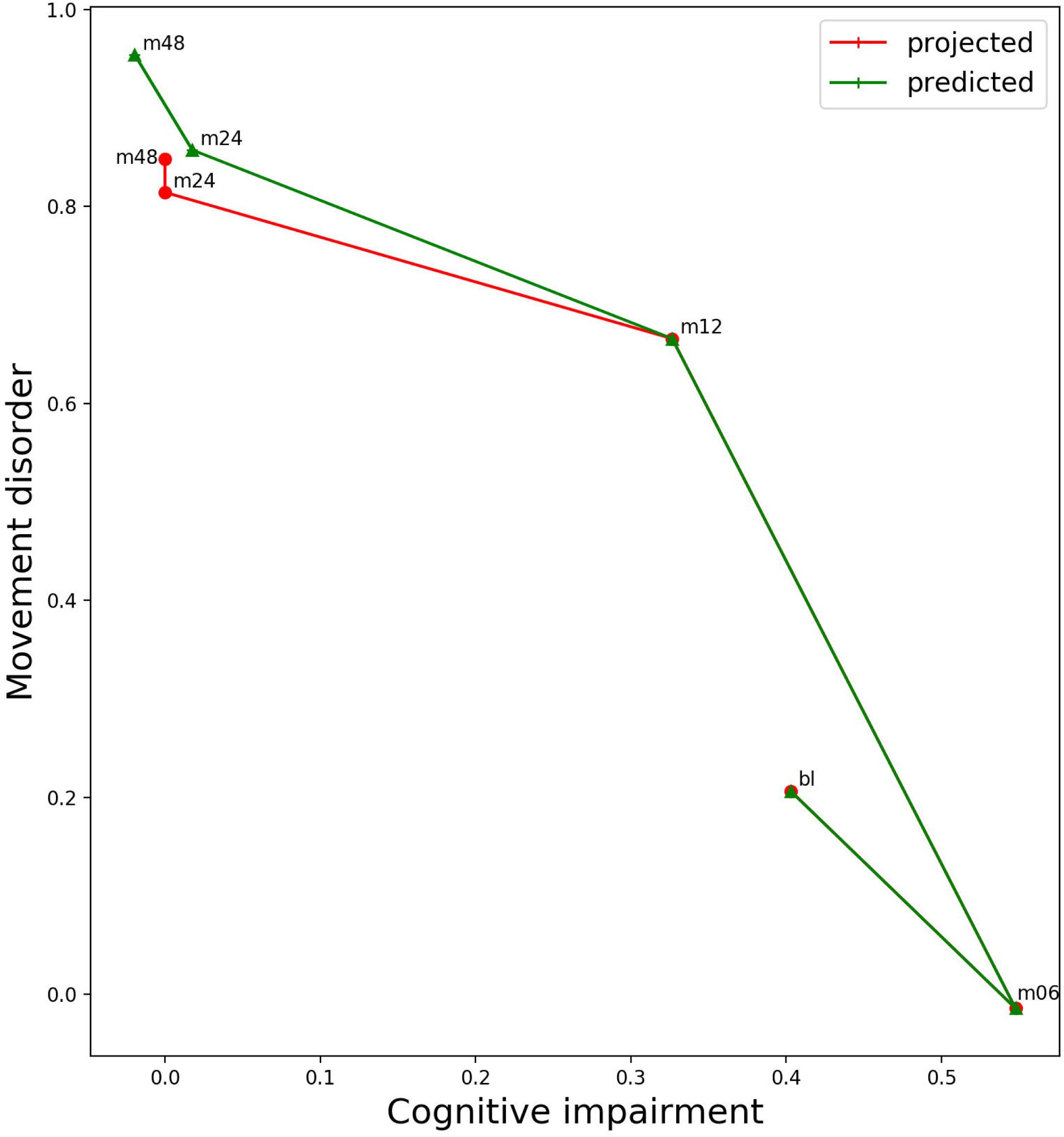
The predicted and projected trajectories of AD progression using LSTM for an AD patient.

NMF was used to project the data in progression space for each visit. The data from the first three visits were used to predict the position of an observation in progression space for the next two visits (visit at 48^th^ and the 24^th^ month). The mean (standard deviation) squared Euclidean distance between projected and predicted test observations in the projection space (quantifying the model performance) for the 48^th^ and the 24^th^ month are 0.00553881(0.00144076) and 0.0046597 (0.000620) respectively. The number of participants in each cohort was limited, and not all the features were ascertained in all three (baseline, after 6 and 12 months) visits. These limitations affected the model performance. Further, we compared the LSTM results to a linear regression model. Using linear regression, the mean (standard deviation) squared Euclidean distance between projected and predicted test observations in the projection space for the 48^th^ and the 24th month 0.0375324 (0.0146328) (6.7 times larger than LSTM) and 0.0235437 (0.0226417) (5 times larger) respectively

### Statistics of each progression subtype

The number of subjects with MCI, dementia, and controls in each subtype is shown in Tables 2A and 2B for prediction at 24- and 48-month respectively. Fig 8 shows their share percentage in each of the subtypes present after 24 months from baseline. As expected, the share of dementia patients is maximum in a fast progression rate. The slow progression rate has no dementia patient and contains only MCI patients. The moderate progression rate subtype is dominated by MCI patients, which covers ∼70% of the observations in the moderate subtype. A similar trend was observed after 48 months from baseline as well, as presented in the supplementary section S8.

**Table 2.** A Number of MCI and dementia patients in each subtype for prediction at m24 using 12 months of data

|  | Slow | Moderate | Fast |
| --- | --- | --- | --- |
| MCI | 114 | 68 | 5 |
| **Dementia** | 0 | 30 | 48 |

**Table 2.** B Number of MCI and dementia patients in each subtype for prediction at m48 using 12 months of data

|  | **Slow** | **Moderate** | **Fast** |
| --- | --- | --- | --- |
| **MCI** | 42 | 36 | 12 |
| **Dementia** | 0 | 2 | 32 |

**Fig 8.**
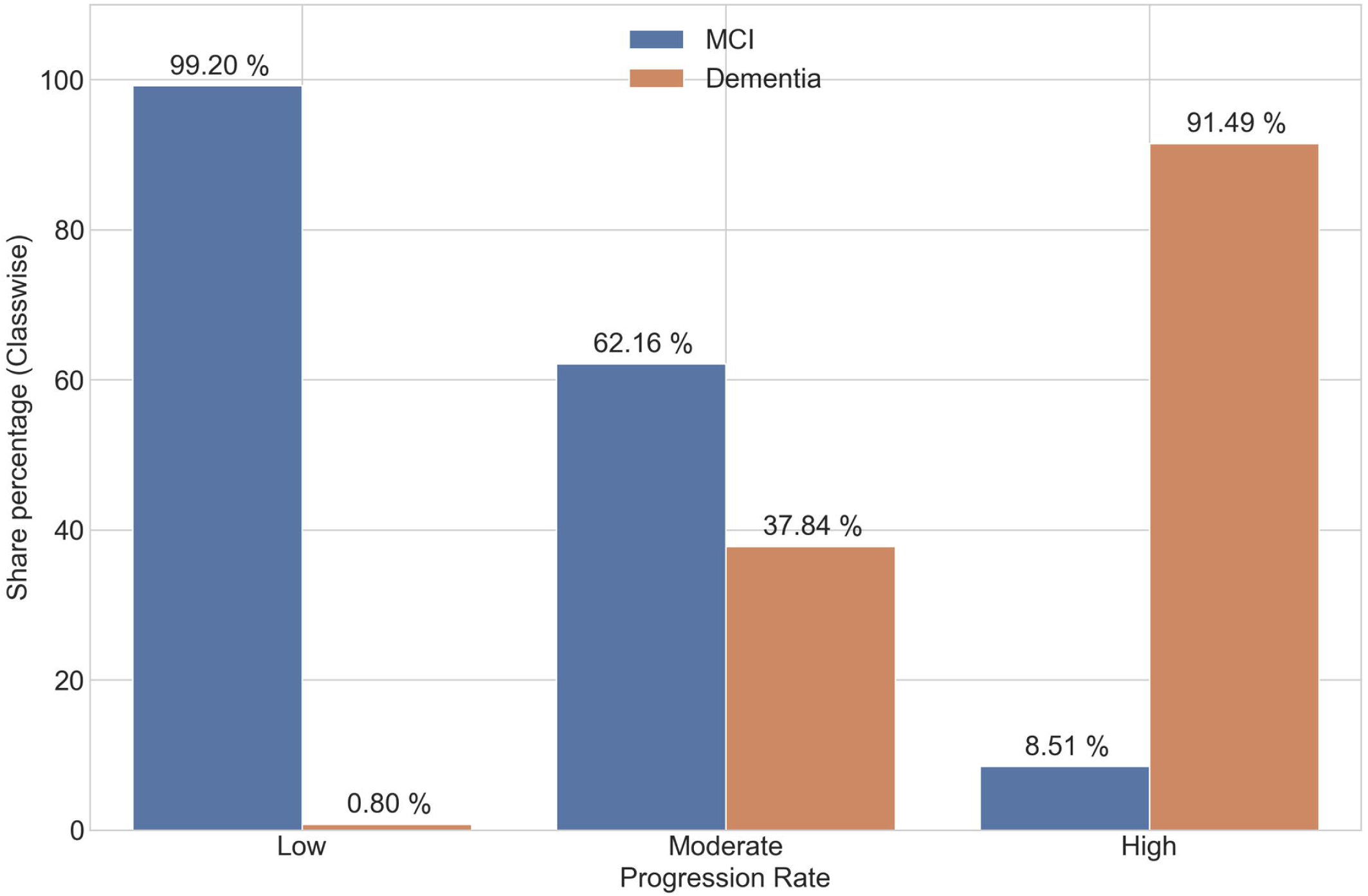
Percent share of controls, MCI, and dementia patients for different subtypes present after 24 months from baseline. The share of MCI patients is decreasing with an increase in progression rate.

### Percent share of APOEε4 variants for different subtypes

APOE has three common alleles, ε2, ε3, and ε4, of which the ε4 allele is closely associated with increased risk of AD [36]. The distribution of APOEε4 alleles for each progression rate subtype is shown in Fig 9 and S9 (in the supplementary material) after 24 and 48 months from the baseline, respectively. This figure illustrates that the progression rate increases with the number of APOEε4 alleles. The absolute counts of 0, 1, and 2 occurrences of APOEε4 alleles in each progression rate are listed in the supplementary section.

**Fig 9.**
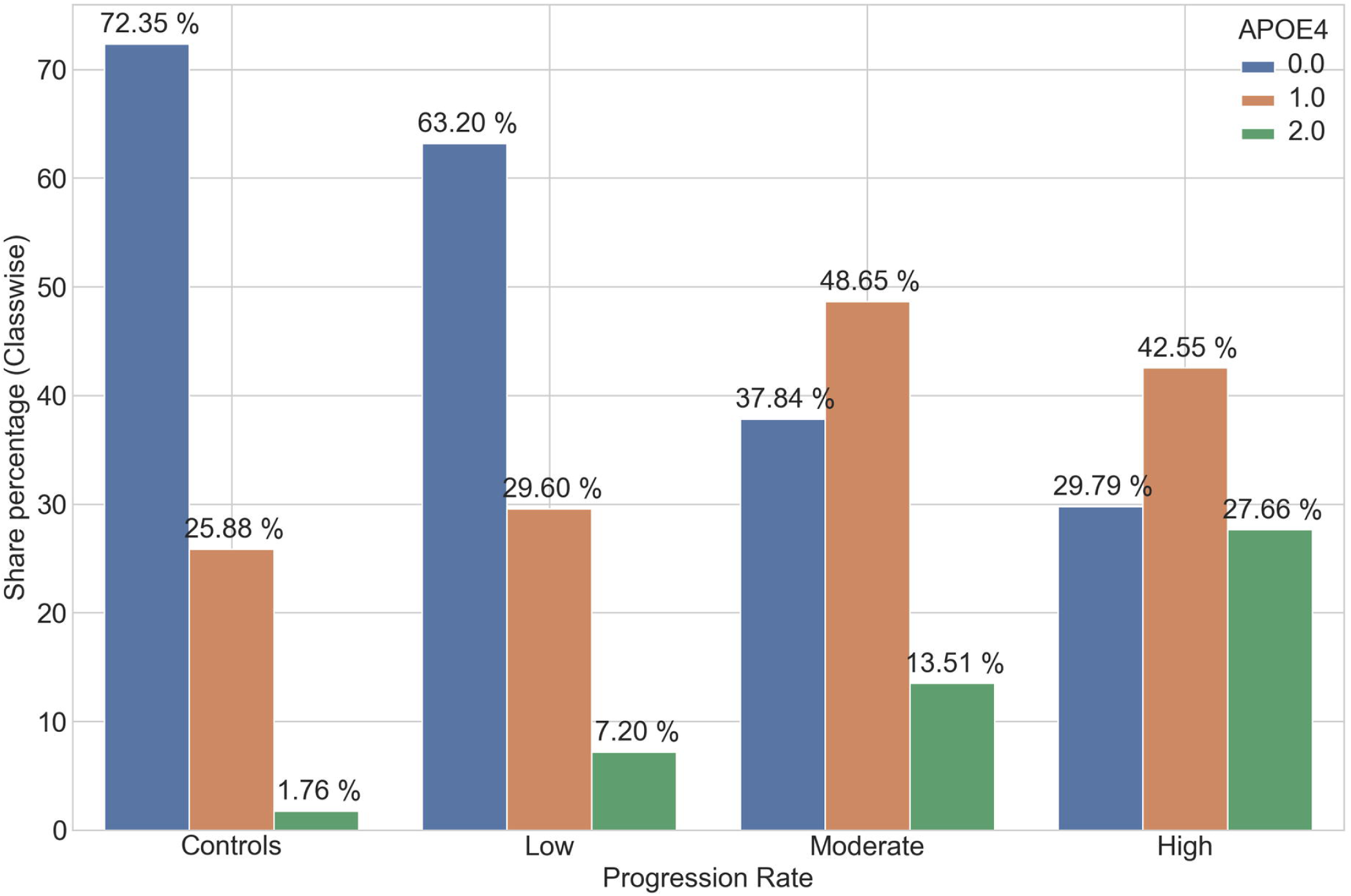
Percent share of APOEε4 alleles for different subtypes after 24 months from baseline. The share of 0 occurrences of APOEε4 alleles is decreasing with an increase in progression rate, whereas the share of 1 and 2 occurrences are increasing.

## Discussion

In this section, we study the reversion of AD captured in the constructed progression space, correlation between the APOEε4 genetic variants and participant’s progression state, effects of aging on AD progression in controls, correlation of memory decline in AD patients with their educational and occupational attainments and distribution of projected dimensions (memory decline and cognitive decline) for each AD subtype, the correlation between certain selective features and AD progression rates in the following subsections. To conclude, we discuss some of the future directions to extend the proposed study.

### AD progression space and disease reversion

Since ADNI is a longitudinal study, the disease state of patients is reassessed every 12 months. The clinical condition either deteriorated or stayed the same for most of the patients but in rare instances, it reversed to a better state, i.e. some patients were observed moving from dementia to MCI or MCI to control stage. These observations were plotted to assess the robustness of the constructed progression space. Fig 10 plots these reversion cases at the 24^th^ and 48^th^ month. It can be verified from these figures that patients moving from dementia to MCI fall in the intermediate region between dementia and MCI (moderate progression rate region). Similarly, patients moving from MCI to control lie in the intermediate progression region between them. Thus, the progression space captures the reversion of the disease state.

**Fig 10.**
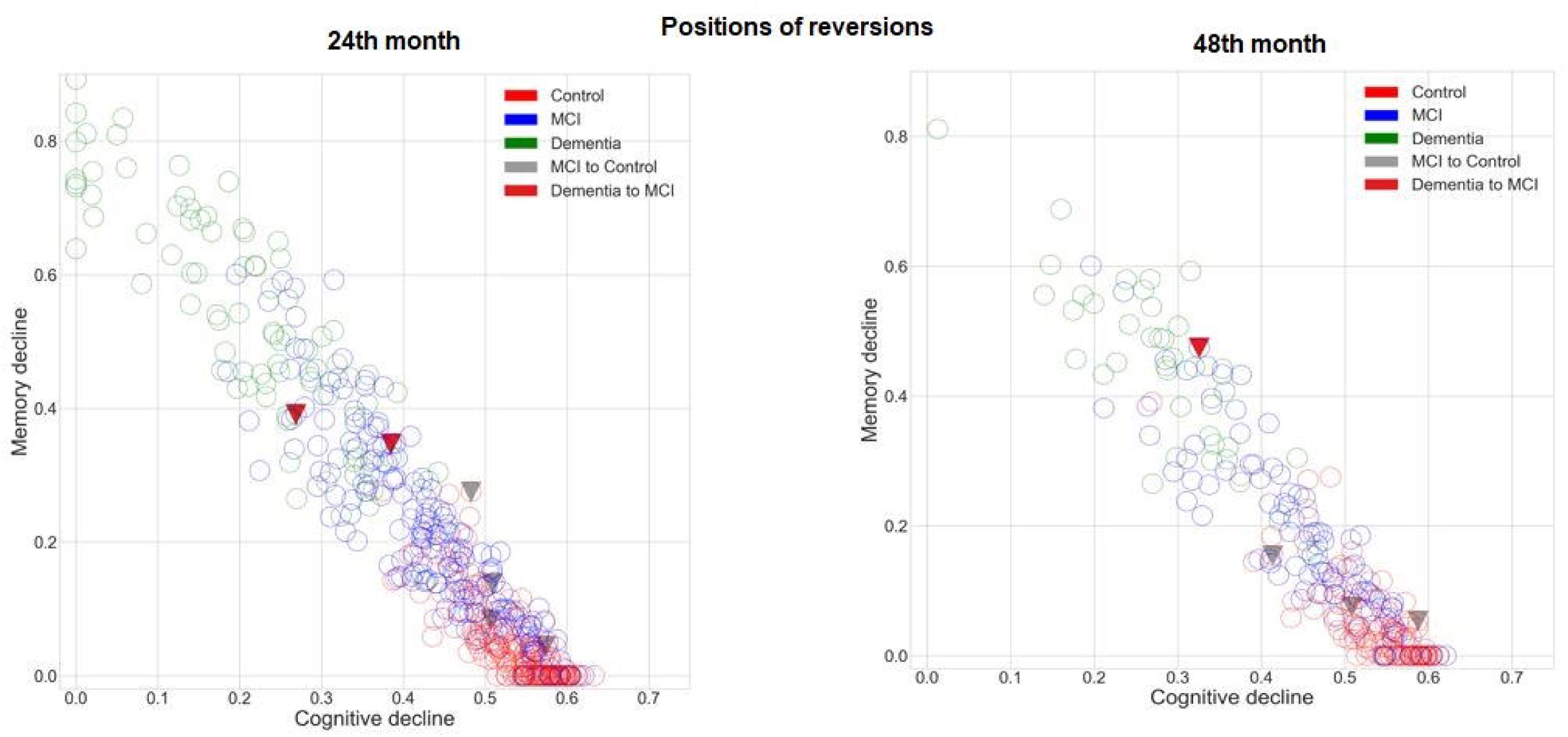
Label Reversions (only MCI to control and dementia to MCI). **Left:** The positions of the reversions at 24^th^ month relative to all other observations. **Right:** The positions of the reversions at 48^th^ month relative to all other observations.

### AD progression states and APOEε4 allele counts

To understand the underlying biological patterns among patients in the progression space, we plotted the distribution of the APOEε4 alleles. Fig 11 projects the observations with 0 and 2 counts of APOEε4 variants on the AD progression space at 24^th^ and 48^th^ month. It is evident from these figures that observations with a 0 count for the APOEε4 allele are concentrated towards the low progression rate zone, whereas observations with 2 counts of APOEε4 allele are concentrated towards the moderate and high progression rate zones. This observation further validates the existing literature [37] and confirms a significant correlation between APOEε4 with cognitive performance.

**Fig 11.**
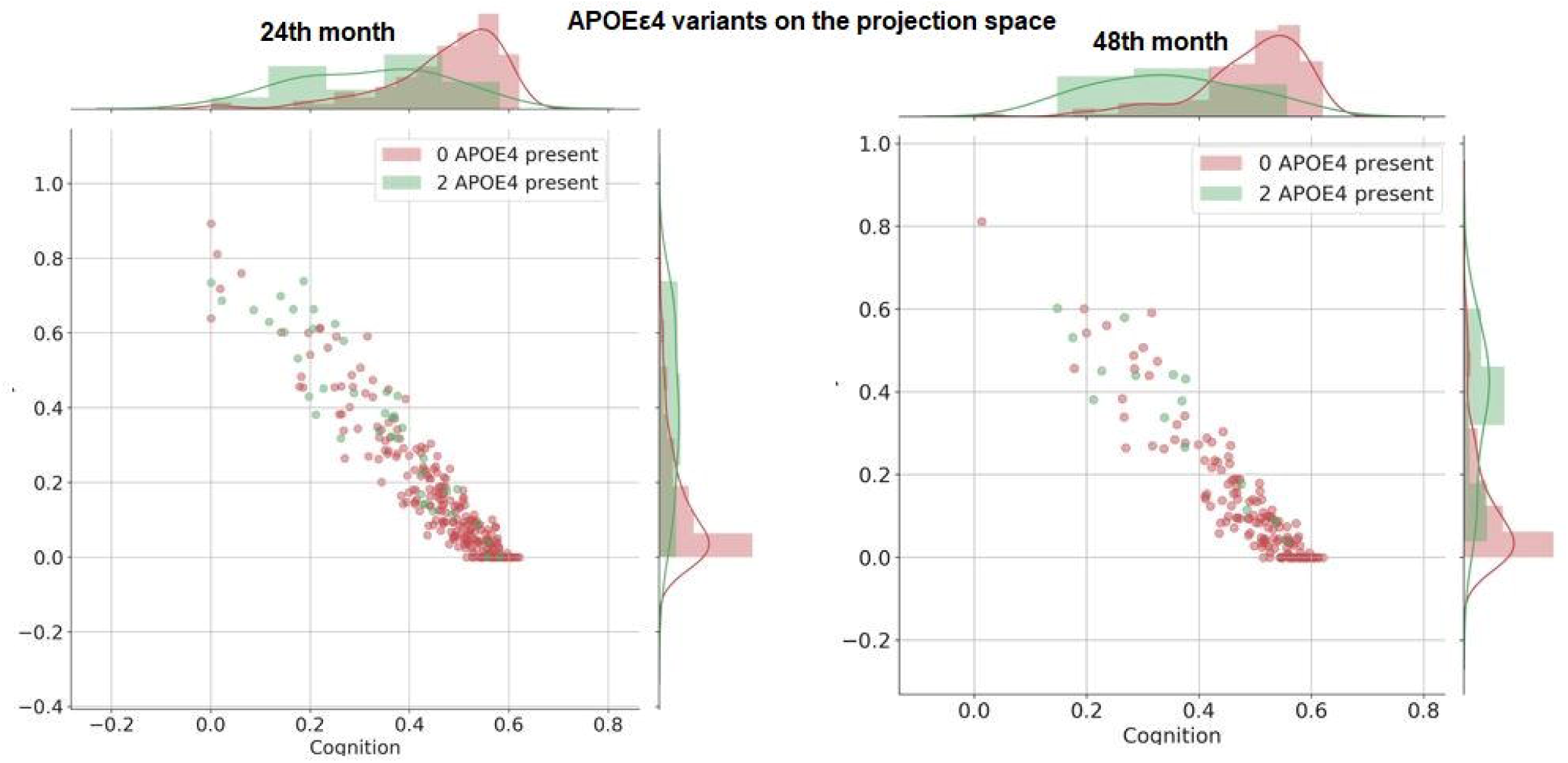
Projection of the number of APOEε4 variants on the projection space. x-axis and y-axis have visualizations of the distribution of APOEε4 alleles in those directions. **Left:** at 24^th^ month. **Right:** at 48^th^ month.

### AD progression in controls and aging

In Fig 12, progression can also be seen in control observations at 24^th^ and 48^th^ month respectively, attributed to a decline in normal cognition and memory with increasing age of the participants. Since this decline is not severe, the observations do not lie in moderate or high progression rate zones. A simple clustering of observations into two clusters shows a stark difference in the mean age of the clusters. It is interesting to note that the mean age for the cluster, which is relatively close to the moderate progression rate zone is 74.59 years and the mean age of the cluster away from this zone is 72.15 years. Similarly, for the 48^th^ month, the mean age of the two clusters relatively close and away from the moderate progression zone are 71.53 and 73.51 respectively (Fig 12).

**Fig 12.**
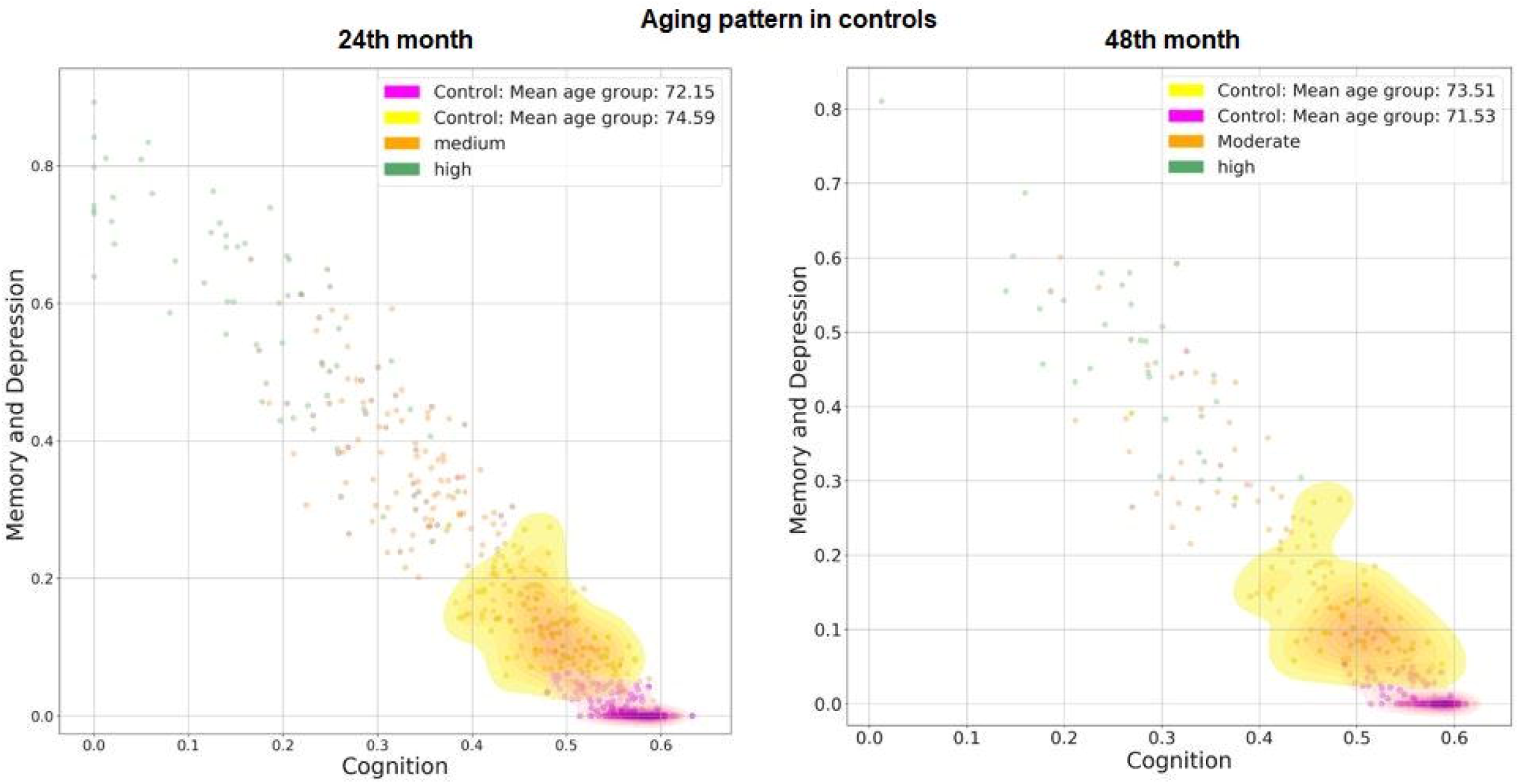
Aging pattern in controls. Mean age of cases in the two clusters of controls. Clusters represent an aging pattern in controls. The cluster near moderate progression rate zone has a high mean age than one which is away from it. **Left:** at 24^th^ month. **Right:** at 48^th^ month.

### Memory decline in AD patients and educational acquirements

To further discover a generalized trend in the AD progression rate, a polynomial curve was fitted on the projected observations. BIC was used to find out the optimum degree of the fitting polynomial, which was observed to be three. As seen in Fig 13 (Left), the cubic curve fits the data in a linear fashion in the low progression region. However, it deviates slightly from this linear behavior in high and moderate progression regions. The magnitude of the slope of the linear curve is 1.19 indicating a rapid memory decline as compared to cognitive. The slope of the progression for 200 most and least educated observations is shown in Fig 13 (Right). The magnitude of slope for the linear curve is 1.26 and 1.19 for highly educated and less educated patients respectively. As the slope for highly educated patients is greater than the slope for less educated patients, it can be inferred that there is a relatively rapid decline in memory of patients with higher education. A study on the links between education and memory decline in AD was carried out in [38]. The research concluded that memory declined more rapidly in AD patients with higher educational and occupational attainment. Thus, our results are further validated by these explorations done in the previous research [38].

**Fig 13.**
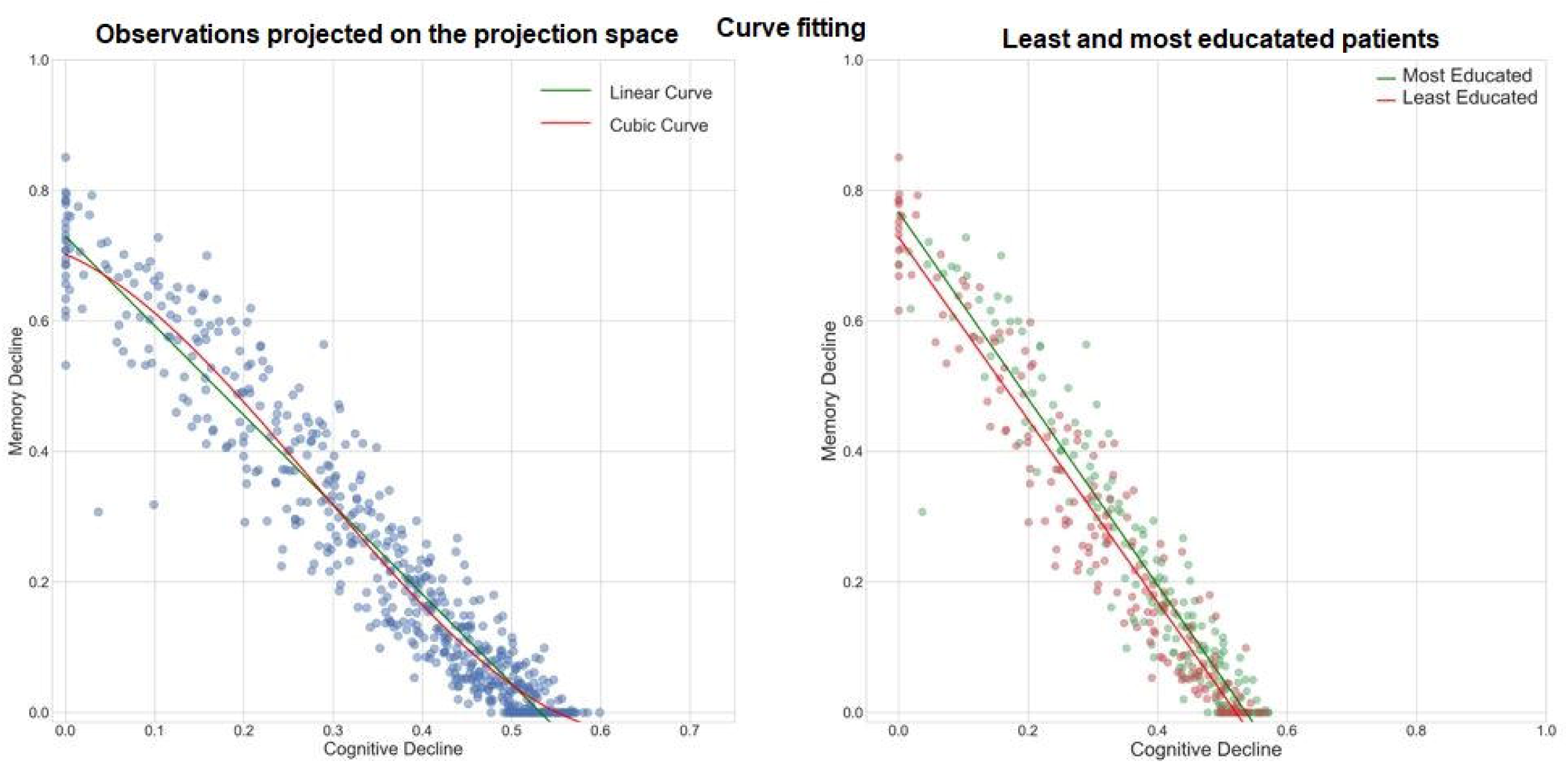
Memory decline in AD patients and educational acquirements. **Left:** Linear and cubic curve-fitting on 582 observations projected on the progression space. The magnitude of the slope of the linear curve is 1.19 indicating a rapid memory decline as compared to cognitive. **Right:** Linear curve fitting for 200 most educated and 200 least educated patients.

### Distribution of memory and cognitive decline

Fig 14 shows the distribution of projected dimensions (memory decline and cognitive decline) for each AD subtype after 24 and 48 months. In the progression space, along the positive direction of the y-axis, the memory decline increases, and along the negative direction of the x-axis the cognitive decline increases. A low value on the x-axis indicates higher cognitive decline whereas a high value on the y-axis indicates higher memory decline. High progression rate has the highest memory and cognitive decline, which goes on reducing with a reduction in progression rate.

**Fig 14.**
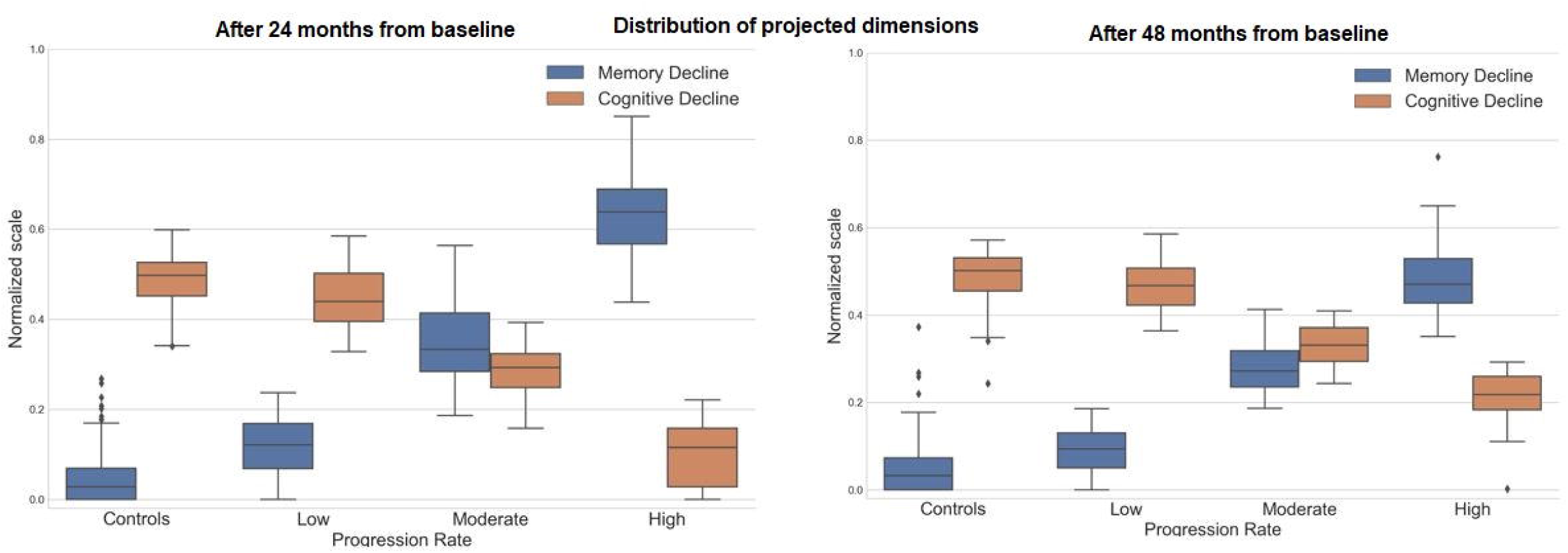
Distribution of projected dimensions (cognitive and memory decline) for each AD progression subtype. A lower numeric value on the cognition axis indicates high cognitive decline whereas a higher numeric value on the memory axis indicates high memory decline. A similar relationship can be observed in the figures. **Left:** after 24 months from baseline. **Right:** after 48 months from baseline.

### Selective features and AD progression rates

Fig 15 shows the distribution of the MMSE score after 6 and 12 months for each AD subtype at the 24^th^ month. Reduction in the MMSE score with increased progression is observed. A similar trend is observed in the distribution of the MMSE score for each AD subtype at the 48^th^ month (given in the supplementary section S10). Further, there is an increase in functional assessment questionnaire (FAQ) total score with increasing progression rate for the 24th and 48^th^ month AD subtypes as shown in Fig 15 and supplementary section S11 respectively.

**Fig 15.**
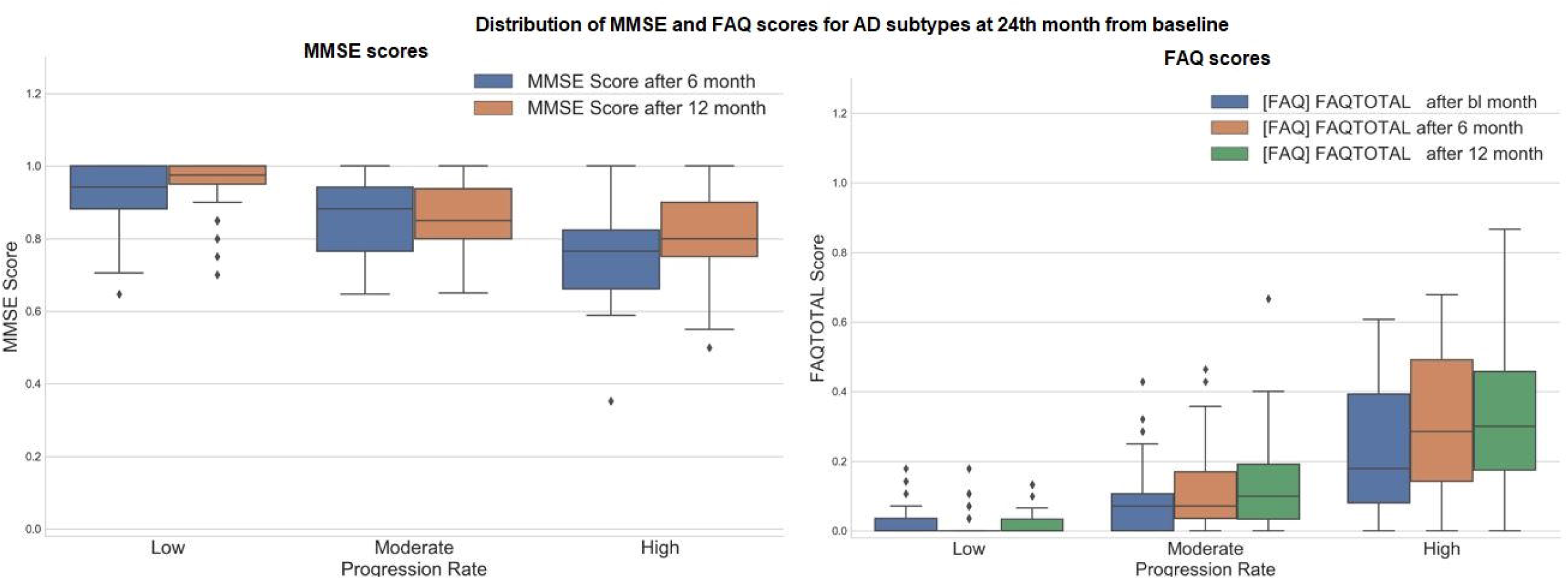
**Left:** Distribution of the normalized MMSE score between 0 and 1 for each AD subtype (subtypes at 24^th^ month). MMSE score decreases with an increase in progression rate. **Right:** Distribution of FAQ total score for each AD subtype (subtypes at 24^th^ month). FAQ total score increases with an increase in progression rate.

### Limitations and future directions

In this work, we discussed the share of different APOEε4 genotype for each progression rate and its correlation with cognitive performance. Future work should involve examining a few other genes that also have been closely related to the progression of AD for studying their interactions [39], [40]. As stated earlier in the paper, AD risk is associated with APOEε4 gene variants [36]. However, the progression space was constructed using only the time-variant clinical data.

Therefore, APOEε4 data were not considered during the construction of the projection space. Moreover, the diagnosis of participants (control, MCI, AD) in the ADNI study is based on their clinical examinations, having a sensitivity of 70.9-87.3% as compared to the neuropathologic assessments, which are considered the gold standard for AD identification [41]. Hence, the discussed progression models suffer from the implicit noise involved in the diagnosis of the study participants. For future analysis, involving diagnosis with neuropathologic examinations may help scale down the ambiguity involved in the true status of participants [40].

The present analysis can be continued in various directions. Since both AD and PD are neurological diseases with AD primarily affecting memory advancing to influence motor functions and PD impacting movement and coordination progressing to hinder memory and other cognitive processes [42], [43], exploring to project ADNI with a PD dataset might explain features responsible for PD progressing to AD or vice versa. Moreover, this study involved investigations only considering the clinical data for exploring AD progressions. Integrating further information such as neuroimaging or biomarker data may augment additional information in our analysis. Since we relied on the CV to gauge the performance of our models, validating the study on separate AD datasets [44], [45]. Finally, the discussed analysis only focused on predicting progression space in AD. The proposed framework can be further adapted to study additional kinds of dementia, such as frontotemporal degenerations, Lewy body dementia, multi-infarct dementia, etc.

## Conclusions

This work clusters participants in distinct progression stages of AD and discusses an approach to predict the future rate of progression after the 24^th^ and 48^th^ months from baseline using longitudinal clinical data. Predicting disease progression serves as a paramount challenge in the therapy and cure of several elaborate diseases. This study is a step forward towards designing sophisticated machine-learning paradigms to facilitate early diagnosis of AD progression. Predicting AD progression rates would lead to better patient-specific attention by recognizing at an early stage the patients with a swift rate of progression. The proposed disease progression and trajectory prediction algorithms can help doctors and practitioners develop a methodical and organized course for clinical tests, which can be much more concise and effective in detection. These adaptations and modifications in clinics may help to diminish treatment and therapy costs for dementia. Further, no healing treatments in AD modification exist, hence, the capability to anticipate the trajectory of impending AD progression at the early stages of the disease is an advancement towards uncovering novel treatments for AD modification. The study provides insights into the progression of AD-related symptoms and signs.

## Acknowledgments

This work was supported in part by the Intramural Research Programs of the National Institute on Aging and the National Institute of Neurological Disorders and Stroke (project numbers: Z01-AG000949-02 and ZIA-NS003154). Data collection and sharing for this project was funded by the Alzheimer’s Disease Neuroimaging Initiative (ADNI) (National Institutes of Health Grant U01 AG024904) and DOD ADNI (Department of Defense award number W81XWH-12-2-0012). ADNI is funded by the National Institute on Aging, the National Institute of Biomedical Imaging and Bioengineering, and through generous contributions from the following: AbbVie, Alzheimer’s Association; Alzheimer’s Drug Discovery Foundation; Araclon Biotech; BioClinica, Inc.; Biogen; Bristol-Myers Squibb Company; CereSpir, Inc.; Cogstate; Eisai Inc.; Elan Pharmaceuticals, Inc.; Eli Lilly and Company; EuroImmun; F. Hoffmann-La Roche Ltd and its affiliated company Genentech, Inc.; Fujirebio; GE Healthcare; IXICO Ltd.; Janssen Alzheimer Immunotherapy Research & Development, LLC.; Johnson & Johnson Pharmaceutical Research & Development LLC.; Lumosity; Lundbeck; Merck & Co., Inc.; Meso Scale Diagnostics, LLC.; NeuroRx Research; Neurotrack Technologies; Novartis Pharmaceuticals Corporation; Pfizer Inc.; Piramal Imaging; Servier; Takeda Pharmaceutical Company; and Transition Therapeutics. The Canadian Institutes of Health Research is providing funds to support ADNI clinical sites in Canada. Private sector contributions are facilitated by the Foundation for the National Institutes of Health (www.fnih.org). The grantee organization is the Northern California Institute for Research and Education, and the study is coordinated by the Alzheimer’s Therapeutic Research Institute at the University of Southern California.

## REFERENCES

1. Corrada MM, Brookmeyer R, Paganini-Hill A, Berlau D, Kawas CH. Dementia incidence continues to increase with age in the oldest old: The 90 study. Ann Neurol. 2010;67: 114–121.

2. Ferri CP, Prince M, Brayne C, Brodaty H, Fratiglioni L, Ganguli M, et al. Global prevalence of dementia: a Delphi consensus study. Lancet. 2005;366: 2112–2117.

3. Villemagne VL, Burnham S, Bourgeat P, Brown B, Ellis KA, Salvado O, et al. Amyloid β deposition, neurodegeneration, and cognitive decline in sporadic Alzheimer’s disease: a prospective cohort study. The Lancet Neurology. 2013. pp. 357–367. doi: 10.1016/s1474-4422(13)70044-9

4. Jack CR Jr, Lowe VJ, Weigand SD, Wiste HJ, Senjem ML, Knopman DS, et al. Serial PIB and MRI in normal, mild cognitive impairment and Alzheimer’s disease: implications for sequence of pathological events in Alzheimer’s disease. Brain. 2009;132: 1355–1365.

5. Reiman EM, Quiroz YT, Fleisher AS, Chen K, Velez-Pardo C, Jimenez-Del-Rio M, et al. Brain imaging and fluid biomarker analysis in young adults at genetic risk for autosomal dominant Alzheimer’s disease in the presenilin 1 E280A kindred: a case-control study. The Lancet Neurology. 2012. pp. 1048–1056. doi: 10.1016/s1474-4422(12)70228-4

6. Bateman RJ, Xiong C, Benzinger TLS, Fagan AM, Goate A, Fox NC, et al. Clinical and biomarker changes in dominantly inherited Alzheimer’s disease. N Engl J Med. 2012;367: 795–804.

7. Braak H, Thal DR, Ghebremedhin E, Del Tredici K. Stages of the pathologic process in Alzheimer disease: age categories from 1 to 100 years. J Neuropathol Exp Neurol. 2011;70: 960–969.

8. Alzheimer’s Association. 2019 alzheimer’s disease facts and figures report. 2019 [cited 11 Sep 2019]. Available: https://www.alz.org/media/documents/alzheimers-facts-and-figures-2019-r.pdf

9. Alzheimer’s Association. Alzheimer’s Disease and Dementia: Medications for Memory Loss. [cited 11 Sep 2019]. Available: https://alz.org/alzheimers-dementia/treatments/medications-for-memory

10. Alzheimer’s Association. Treatments for Alzheimer’s Disease and Dementia. [cited 11 Sep 2019]. Available: https://alz.org/alzheimers-dementia/treatments

11. Alzheimer’s Association. Alzheimer’s Disease and Dementia: Stages of Alzheimer’s. [cited 12 Sep 2019]. Available: https://alz.org/alzheimers-dementia/stages

12. Holtzman DM, Morris JC, Goate AM. Alzheimer’s Disease: The Challenge of the Second Century. Science Translational Medicine. 2011. pp. 77sr1–77sr1. doi: 10.1126/scitranslmed.3002369

13. Boyle PA, Wilson RS, Aggarwal NT, Tang Y, Bennett DA. Mild cognitive impairment: risk of Alzheimer disease and rate of cognitive decline. Neurology. 2006;67: 441–445.

14. Hansson O, Zetterberg H, Buchhave P, Londos E, Blennow K, Minthon L. Association between CSF biomarkers and incipient Alzheimer’s disease in patients with mild cognitive impairment: a follow-up study. Lancet Neurol. 2006;5: 228–234.

15. Asif S, Khan T. A Machine Learning Model to Predict the Onset of Alzheimer Disease using Potential Cerebrospinal Fluid (CSF) Biomarkers. International Journal of Advanced Computer Science and Applications. 2017;8. doi: 10.14569/ijacsa.2017.081216

16. Moreland J, Urhemaa T, van Gils M, Lötjönen J, Wolber J, Buckley CJ. Validation of prognostic biomarker scores for predicting progression of dementia in patients with amnestic mild cognitive impairment. Nucl Med Commun. 2018;39: 297–303.

17. Altaf T, Anwar SM, Gul N, Majeed MN, Majid M. Multi-class Alzheimer’s disease classification using image and clinical features. Biomed Signal Process Control. 2018;43: 64–74.

18. Faghri F, Hashemi SH, Leonard H, Scholz SW, Campbell RH, Nalls MA, et al. Predicting onset, progression, and clinical subtypes of Parkinson disease using machine learning. 2018. doi: 10.1101/338913

19. Satone V, Kaur R, Faghri F, Nalls MA, Singleton AB, Campbell RH. Learning the progression and clinical subtypes of Alzheimer’s disease from longitudinal clinical data. arXiv [cs.LG]. 2018. Available: http://arxiv.org/abs/1812.00546

20. ADNI Procedures Manuals. [cited 7 Sep 2019]. Available: https://adni.loni.usc.edu/wp-content/uploads/2010/09/ADNI_GeneralProceduresManual.pdf

21. Nasreddine ZS, Phillips NA, BÃ©dirian V, Charbonneau S, Whitehead V, Collin I, et al. The Montreal Cognitive Assessment, MoCA: A Brief Screening Tool For Mild Cognitive Impairment. Journal of the American Geriatrics Society. 2005. pp. 695–699. doi: 10.1111/j.1532-5415.2005.53221.x

22. Berg L. Clinical Dementia Rating. British Journal of Psychiatry. 1988;153: 267.

23. Zaidi S, Kat MG, de Jonghe JFM. Clinician and caregiver agreement on neuropsychiatric symptom severity: a study using the Neuropsychiatric Inventory – Clinician rating scale (NPI-C). International Psychogeriatrics. 2014. pp. 1139–1145. doi: 10.1017/s1041610214000295

24. Dodrill CB. A Neuropsychological Battery for Epilepsy. Epilepsia. 1978;19: 611–623.

25. Folstein MF, Folstein SE, McHugh PR. Mini-mental State: A Practical Method for Grading the Cognitive State of Patients for the Clinician. 1975.

26. Yesavage JA, Brink TL, Rose TL, Lum O, Huang V, Adey M, et al. Geriatric Depression Scale. PsycTESTS Dataset. 1983. doi: 10.1037/t00930-000

27. Allaire JC, Marsiske M. Everyday cognition: Age and intellectual ability correlates. Psychol Aging. 1999;14: 627–644.

28. Salomon G, Perkins DN, Globerson T. Partners in Cognition: Extending Human Intelligence with Intelligent Technologies. Educ Res. 1991;20: 2.

29. Fillenbaum GG, Smyer MA. The development, validity, and reliability of the OARS multidimensional functional assessment questionnaire. J Gerontol. 1981;36: 428–434.

30. Rosen WG, Mohs RC, Davis KL. A new rating scale for Alzheimer’s disease. Am J Psychiatry. 1984;141: 1356–1364.

31. Mohs RC, Knopman D, Petersen RC, Ferris SH, Ernesto C, Grundman M, et al. Development of cognitive instruments for use in clinical trials of antidementia drugs: additions to the Alzheimer’s Disease Assessment Scale that broaden its scope. The Alzheimer’s Disease Cooperative Study. Alzheimer Dis Assoc Disord. 1997;11 Suppl 2: S13–21.

32. Lee DD, Seung HS. Algorithms for Non-negative Matrix Factorization. In: Leen TK, Dietterich TG, Tresp V, editors. Advances in Neural Information Processing Systems 13. MIT Press; 2001. pp. 556–562.

33. McLachlan GJ, Basford KE. Mixture models: Inference and applications to clustering. M. Dekker New York; 1988.

34. Schwarz G. Estimating the Dimension of a Model. Ann Stat. 1978;6: 461–464.

35. Breiman L. Random Forests. Mach Learn. 2001;45: 5–32.

36. Strittmatter WJ, Weisgraber KH, Huang DY, Dong LM, Salvesen GS, Pericak-Vance M, et al. Binding of human apolipoprotein E to synthetic amyloid beta peptide: isoform-specific effects and implications for late-onset Alzheimer disease. Proc Natl Acad Sci U S A. 1993;90: 8098–8102.

37. Raber J, Wong D, Yu G-Q, Buttini M, Mahley RW, Pitas RE, et al. Apolipoprotein E and cognitive performance. Nature. 2000;404: 352–354.

38. Stern Y E al. Rate of memory decline in AD is related to education and occupation: cognitive reserve? - PubMed - NCBI. [cited 24 Nov 2018]. Available: https://www.ncbi.nlm.nih.gov/pubmed/10599762

39. Lambert JC, Ibrahim-Verbaas CA, Harold D, Naj AC, Sims R, Bellenguez C, et al. Meta-analysis of 74,046 individuals identifies 11 new susceptibility loci for Alzheimer’s disease. Nat Genet. 2013;45: 1452–1458.

40. Korolev IO, Symonds LL, Bozoki AC, Alzheimer’s Disease Neuroimaging Initiative. Predicting Progression from Mild Cognitive Impairment to Alzheimer’s Dementia Using Clinical, MRI, and Plasma Biomarkers via Probabilistic Pattern Classification. PLoS One. 2016;11: e0138866.

41. Beach TG, Monsell SE, Phillips LE, Kukull W. Accuracy of the clinical diagnosis of Alzheimer disease at National Institute on Aging Alzheimer Disease Centers, 2005-2010. J Neuropathol Exp Neurol. 2012;71: 266–273.

42. Is Alzheimer’s disease related to Parkinson’s disease? | Alzheimer’s Disease. In: Sharecare [Internet]. [cited 8 Sep 2019]. Available: https://www.sharecare.com/health/alzheimers-disease/alzheimers-disease-related-parkinsons-disease

43. Website. [cited 8 Sep 2019]. Available: http://journals.plos.org/plosmedicine/article/file?id=10.1371/journal.pmed.1002266$\&$type=printable

44. AddNeuroMed. In: Consortiapedia [Internet]. [cited 9 Sep 2019]. Available: https://consortiapedia.fastercures.org/consortia/anm/

45. Beekly DL, Ramos EM, Lee WW, Deitrich WD, Jacka ME, Wu J, et al. The National Alzheimer??s Coordinating Center (NACC) Database: The Uniform Data Set. Alzheimer Disease & Associated Disorders. 2007. pp. 249–258. doi: 10.1097/wad.0b013e318142774e

