## supplementary for "Predicting Alzheimer’s disease progression trajectory and clinical subtypes using machine learning"

**Supporting Information**

In this additional section, we highlight some of the descriptive information and statistics supporting our proposed analysis.

**S1: Key eligibility criteria for ADNI participants**

Enrolled participants for the study were between 55-90 (inclusive) years of age, had a study partner, were able to provide an independent evaluation of functioning and were able to speak either English or Spanish. All participants were willing and able to undergo all test procedures including neuroimaging and agreed to longitudinal follow up. Normal, MCI and mild AD participants had MMSE scores between 24-30 (inclusive), 24-30 (inclusive) and 20-26 (inclusive) respectively and a CDR total scores of 0, 0.5 and 1.0 respectively. Refer [[20]](https://paperpile.com/c/4aT5Jb/YPL1) for further details on protocols and criteria for the participants.

**S2 Table: Description of ADNI clinical features used in analysis**

| **Clinical test** | **Sub Test name** | **Features** | **Description** |
| --- | --- | --- | --- |
| Montreal Cognitive Assessment |  |  |  |
| Moca (856, 42) - ['bl', 'm06', m12'] | ALTERNATING TRAIL MAKING | TRAILS |  |
|  | VISUOCONSTRUCTIONAL SKILLS | CUBE | Copy Cube |
|  |  | CLOCKCON | Draw Clock - Contour |
|  |  | CLOCKNO | Draw Clock - Number |
|  |  | CLOCKHAN | Draw Clock - Hands |
|  | NAMING | LION | Lion |
|  |  | RHINO | Rhinoceros |
|  |  | CAMEL | Camel |
|  | MEMORY | IMMT2W1 | Immediate (#2): Daisy |
|  |  | IMMT2W2 | Immediate (#2): Red |
|  |  | IMMT2W3 | Immediate (#2): Velvet |
|  |  | IMMT2W4 | Immediate (#2): Church |
|  |  | IMMT2W5 | Immediate (#1): Daisy |
|  |  | IMMT1W1 | Immediate (#1): Red |
|  |  | IMMT1W2 | Immediate (#2): Face |
|  |  | IMMT1W3 | Immediate (#1): Velvet |
|  |  | IMMT1W4 | Immediate (#1): Church |
|  |  | IMMT1W5 | Immediate (#1): Face |
|  | SENTENCE REPETITION | REPEAT1 | Repeat Sentence: I only know that John is the one to help today. |
|  |  | REPEAT2 | Repeat Sentence: The cat always hid under the couch when dogs were in the room. |
|  | VERBAL FLUENCY | FFLUENCY | Letter Fluency - F: Total number of correct words |
|  | ABSTRACTION | ABSTRAN | Abstraction: train-bicycle |
|  |  | ABSMEAS | Abstraction: watch-ruler |
|  | DELAYED RECALL | DELW1 | Delayed: Face |
|  |  | DELW2 | Delayed: Velvet |
|  |  | DELW3 | Delayed: Church |
|  |  | DELW4 | Delayed: Daisy |
|  |  | DELW5 | Delayed: Red |
|  | ORIENTATION | DATE | - |
|  |  | MONTH | - |
|  |  | YEAR | - |
|  |  | DAY | - |
|  |  | PLACE | - |
|  |  | CITY | - |
|  | ATTENTION | DIGFOR | Digits Forward |
|  |  | DIGBACK | Digits Backward |
|  |  | LETTERS | List of Letters/Tapping: # Errors |
|  |  | SERIAL1 | 1st Subtraction |
|  |  | SERIAL2 | 2nd Subtraction |
|  |  | SERIAL3 | 3rd Subtraction |
|  |  | SERIAL4 | 4thSubtraction |
|  |  | SERIAL5 | 5th Subtraction |
| Clinical Dementia Rating |  |  |  |
| CDR (1461, 6) - ['bl', 'm06', 'm12'] | CDR RATER to compute the Global CDR Score availabe at <http://www.biostat.wustl.edu/adrc/> | CDMEMORY | Memory Score |
|  |  | CDORIENT | Orientation Score |
|  |  | CDJUDGE | Judgment and Problem Solving Score |
|  |  | CDCOMMUN | Community Affairs Score |
|  |  | CDHOME | Home and Hobbies Score |
|  |  | CDCARE | Personal Care Score |
| Neuropsychiatric Inventory Questionnaire | NPI Score = Frequency x Severity |  |  |
| NPIQ (778, 24) - ['bl', 'm06', 'm12'] | Delusions | NPIA | Does {P} believe that others are stealing from him/her, or planning to harm him/her in some way? |
|  |  | NPIASEV | Severity Ratings for Delusions |
|  | Hallucinations | NPIB | Does {P} act as if he/she hears voices? Does he/she talk to people who are not there? |
|  |  | NPIBSEV | Severity Ratings for Hallucinations |
|  | Agitation/Aggression | NPIC | Is {P} stubborn and resistive to help from others? |
|  |  | NPICSEV | Severity Ratings for. Agitation/Aggression |
|  | Depression/Dysphoria | NPID | Does {P} act as if he/she is sad or in low spirits? Does he/she cry? |
|  |  | NPIDSEV | Severity Ratings for Depression/Dysphoria |
|  | Anxiety | NPIE | Does {P} become upset when separated from you? Does he/she have any other signs of nervousness, such as shortness of breath, sighing, being unable to relax, or feeling excessively tense? |
|  |  | NPIESEV | Severity Ratings for Anxiety |
|  | Elation/Euphoria | NPIF | Does {P} appear to feel too good or act excessively happy? |
|  |  | NPIFSEV | Severity Ratings for Elation/Euphoria |
|  | Apathy/Indifference | NPIG | Does {P} seem less interested in his/her usual activities and in the activities and plans of others? |
|  |  | NPIGSEV | Severity Ratings for Apathy/Indifference |
|  | Disinhibition | NPIH | Does {P} seem to act impulsively? For example, does {P} talk to strangers as if he/she knows them, or does {P} say things that may hurt people's feelings? |
|  |  | NPIHSEV | Severity Ratings for Disinhibition |
|  | Irritability/Lability | NPII | Is {P} impatient or cranky? Does he/she have difficulty coping with delays or waiting for planned activities? |
|  |  | NPIISEV | Severity Ratings for Irritability/Lability |
|  | Aberrant Motor Behavior | NPIJ | Does {P} engage in repetitive activities, such as pacing around the house, handling buttons, wrapping strings, or doing other things repeatedly? |
|  |  | NPIJSEV | Severity Ratings for Aberrant Motor Behavior |
|  | Sleep | NPIK | Does {P} awaken you during the night, rise too early in the morning, or take excessive naps during the day? |
|  |  | NPIKSEV | Severity Ratings for Sleep |
|  | Appetite | NPIL | Has {P} lost or gained weight, or had a change in the food he/she likes? |
|  |  | NPILSEV | Severity Ratings for Appetite |
| Neuropsychological Battery |  |  |  |
| NEUROBAT - (1456, 43) | CLOCK - (1637, 1) - ['bl', 'm06', 'm12'] | CLOCKCIRC | 1. Approximately circular face |
|  |  | CLOCKSYM | 2. Symmetry of number placement |
|  |  | CLOCKNUM | 3. Correctness of numbers |
|  |  | CLOCKHAND | 4. Presence of the two hands |
|  |  | CLOCKTIME | 5. Presence of the two hands, set to ten after eleven |
|  | We consider only the Total Score | CLOCKSCOR | Total Score |
|  | COPY - (1637, 1) - ['bl', 'm06', 'm12'] | COPYCIRC | 1. Approximately circular face |
|  |  | COPYSYM | 2. Symmetry of number placement |
|  |  | COPYNUM | 3. Correctness of numbers |
|  |  | COPYHAND | 4. Presence of the two hands |
|  |  | COPYTIME | 5. Presence of the two hands, set to ten after eleven |
|  | We consider only the Total Score | COPYSCOR | Total Score |
|  | STORY - (1476, 1) - ['m12'] | LIMMTOTAL | Total Number of Story Units Recalled |
|  | Category fluency - Animal examples - (1636, 6) - ['bl', 'm06', 'm12'] | CATANIMSC | Category Fluency (Animals) - Total Correct |
|  |  | CATANPERS | Category Fluency (Animals) - Perseverations |
|  |  | CATANINTR | Category Fluency (Animals) - Intrusions |
|  |  | CATVEGESC | Category Fluency (Vegetables) - Total Correct |
|  |  | CATVGPERS | Category Fluency (Vegetables) - Perseverations |
|  |  | CATVGINTR | Category Fluency (Vegetables) - Intrusions |
|  | Trail Making - (1614, 6) - ['bl', 'm06', 'm12'] | TRAASCOR | Part B - Time to complete |
|  |  | TRAAERRCOM | Errors of Commission |
|  |  | TRAAERROM | Errors of Omission |
|  |  | TRABSCOR | Part A - Time to Complete |
|  |  | TRABERRCOM | Errors of Commission |
|  |  | TRABERROM | Errors of Omission |
|  | AV - (1637, 18) - ['bl', 'm06', 'm12'] | AVDEL30MIN | 30 Minute Delay Total |
|  |  | AVDELERR1 | Total Intrusions |
|  |  | AVDELERR2 | Total Intrusions |
|  |  | AVDELTOT | Recognition Score |
|  |  | AVERR1 | Total Intrusions |
|  |  | AVERR2 | Total Intrusions |
|  |  | AVERR3 | Total Intrusions |
|  |  | AVERR4 | Total Intrusions |
|  |  | AVERR5 | Total Intrusions |
|  |  | AVERR6 | Total Intrusions |
|  |  | AVERRB | Total Intrusions |
|  |  | AVTOT1 | Trial 1 Total |
|  |  | AVTOT2 | Trial 2 Total |
|  |  | AVTOT3 | Trial 3 Total |
|  |  | AVTOT4 | Trial 4 Total |
|  |  | AVTOT5 | Trial 5 Total |
|  |  | AVTOT6 | Trial 6 Total |
|  |  | AVTOTB | List B Total |
|  | Digit score - (785, 1) - ['bl', 'm06', 'm12'] | DIGITSCOR | Total Correct |
|  | Logical memeory test - (1474, 2) - ['m12'] | LDELTOTAL | Total Number of Story Units Recalled |
|  |  | LDELCUE | Reminder given? |
|  | Boston naming test - (1631, 5) - ['bl', 'm06', 'm12'] | BNTSPONT | 1. Total correct without a cue |
|  |  | BNTSTIM | 2. Total semantic cues given |
|  |  | BNTCSTIM | 3. Total correct with a semantic cue |
|  |  | BNTPHON | 4. Total phonemic cues given |
|  |  | BNTCPHON | 5. Number of correct responses following a phonemic cue |
| Mini Mental State Exam |  |  |  |
| MMSE - (1451, 31) - ['m06', 'm12'] |  | MMSCORE | MMSE TOTAL SCORE |
|  |  | MMDATE | 1. What is today's date? |
|  |  | MMYEAR | 2. What is the year? |
|  |  | MMMONTH | 3. What is the month? |
|  |  | MMDAY | 4. What day of the week is today? |
|  |  | MMSEASON | 5. What season is it? |
|  |  | MMHOSPIT | 6. What is the name of this hospital (clinic, place)? |
|  |  | MMFLOOR | 7. What floor are we on? |
|  |  | MMCITY | 8. What town or city are we in? |
|  |  | MMAREA | 9. What county (district, borough, area) are we in? |
|  |  | MMSTATE | 10. What state are we in? |
|  |  | MMBALL | 11. Ball |
|  |  | MMFLAG | 12. Flag |
|  |  | MMTREE | 13. Tree |
|  |  | MMD | 14. D |
|  |  | MML | 15. L |
|  |  | MMR | 16. R |
|  |  | MMO | 17. O |
|  |  | MMW | 18. W |
|  |  | MMBALLDL | 19. Ball |
|  |  | MMFLAGDL | 20. Flag |
|  |  | MMTREEDL | 21. Tree |
|  |  | MMWATCH | 22. Show the participant a wrist watch and ask "What is this?" |
|  |  | MMPENCIL | 23. Repeat for pencil. |
|  |  | MMREPEAT | 24. Say, "Repeat after me: no ifs, ands, or buts." |
|  |  | MMHAND | 25. Takes paper in right hand. |
|  |  | MMFOLD | 26. Folds paper in half. |
|  |  | MMONFLR | 27. Puts paper on floor. |
|  |  | MMREAD | 28. Present the piece of paper which reads, "CLOSE YOUR EYES," and say: "Read this and do what it says." |
|  |  | MMWRITE | 29. Give the participant a blank piece of paper and say: "Write a sentence." |
|  |  | MMDRAW | 30. Present the participant with the Construction Stimulus page. Say, "Copy this design." |
| Geriatric depression scale |  |  |  |
| GDSCALE - (723, 1) - ['m06', 'm12'] |  | GDSATIS | 1. Are you basically satisfied with your life? |
|  |  | GDDROP | 2. Have you dropped many of your activities and interests? |
|  |  | GDEMPTY | 3. Do you feel that your life is empty? |
|  |  | GDBORED | 4. Do you often get bored? |
|  |  | GDSPIRIT | 5. Are you in good spirits most of the time? |
|  |  | GDAFRAID | 6. Are you afraid that something bad is going to happen to you? |
|  |  | GDHAPPY | 7. Do you feel happy most of the time? |
|  |  | GDHELP | 8. Do you often feel helpless? |
|  |  | GDHOME | 9. Do you prefer to stay at home, rather than going out and doing new things? |
|  |  | GDMEMORY | 10. Do you feel you have more problems with memory than most? |
|  |  | GDALIVE | 11. Do you think its wonderful to be alive now? |
|  |  | GDWORTH | 12. Do you feel pretty worthless the way you are now? |
|  |  | GDENERGY | 13. Do you feel full of energy? |
|  |  | GDHOPE | 14. Do you feel that your situation is hopeless? |
|  |  | GDBETTER | 15. Do you think that most people are better off than you are? |
|  | We consider only the Total Score | GDTOTAL | Total Score |
|  |  | ADNI_EF | Executive function summary score |
| Baseline symptoms |  |  |  |
| BLSCHECK - (821, 27) - ['bl'] |  | BCNAUSEA | 1. Nausea |
|  |  | BCVOMIT | 2. Vomiting |
|  |  | BCDIARRH | 3. Diarrhea |
|  |  | BCCONSTP | 4. Constipation |
|  |  | BCABDOMN | 5. Abdominal discomfort |
|  |  | BCSWEATN | 6. Sweating |
|  |  | BCDIZZY | 7. Dizziness |
|  |  | BCENERGY | 8. Low energy |
|  |  | BCDROWSY | 9. Drowsiness |
|  |  | BCVISION | 10. Blurred vision |
|  |  | BCHDACHE | 11. Headache |
|  |  | BCDRYMTH | 12. Dry mouth |
|  |  | BCBREATH | 13. Shortness of breath |
|  |  | BCCOUGH | 14. Coughing |
|  |  | BCPALPIT | 15. Palpitations |
|  |  | BCCHEST | 16. Chest pain |
|  |  | BCURNDIS | 17. Urinary discomfort (e.g., burning) |
|  |  | BCURNFRQ | 18. Urinary frequency |
|  |  | BCANKLE | 19. Ankle swelling |
|  |  | BCMUSCLE | 20. Musculoskeletal pain |
|  |  | BCRASH | 21. Rash |
|  |  | BCINSOMN | 22. Insomnia |
|  |  | BCDPMOOD | 23. Depressed mood |
|  |  | BCCRYING | 24. Crying |
|  |  | BCELMOOD | 25. Elevated mood |
|  |  | BCWANDER | 26. Wandering |
|  |  | BCFALL | 27. Fall |
| Neuropsychiatric Inventory |  |  |  |
| NPI - (627, 12) - ['bl', 'm12'] |  | NPIATOT | A. Delusions: Item score |
|  |  | NPIBTOT | B. Hallucinations: Item score |
|  |  | NPICTOT | C. Agitation/Aggression: Item score |
|  |  | NPIDTOT | D. Depression/Dysphoria: Item score |
|  |  | NPIETOT | E. Anxiety: Item score |
|  |  | NPIFTOT | F. Elation/Euphoria: Item score |
|  |  | NPIGTOT | G. Apathy/Indifference: Item score |
|  |  | NPIHTOT | H. Disinhibition: Item score |
|  |  | NPIITOT | I. Irritability/Lability: Item score |
|  |  | NPIJTOT | J. Aberrant Motor Behavior: Item score |
|  |  | NPIKTOT | K. Sleep: Item score |
|  |  | NPILTOT | L. Appetite and eating disorders: Item score |
| Everyday cognition - study partner |  |  |  |
| ECOGSP - (842, 39) - ['bl', 'm06', 'm12'] |  | MEMORY1 | 1. Remembering a few shopping items without a list. |
|  |  | MEMORY2 | 2. Remembering things that happened recently (such as recent outings, events in the news). |
|  |  | MEMORY3 | 3. Recalling conversations a few days later. |
|  |  | MEMORY4 | 4. Remembering where he/she has placed objects. |
|  |  | MEMORY5 | 5. Repeating stories and/or questions. |
|  |  | MEMORY6 | 6. Remembering the current date or day of the week. |
|  |  | MEMORY7 | 7. Remembering he/she has already told someone something. |
|  |  | MEMORY8 | 8. Remembering appointments, meetings, or engagements. |
|  |  | LANG1 | 1. Forgetting the names of objects. |
|  |  | LANG2 | 2. Verbally giving instructions to others. |
|  |  | LANG3 | 3. Finding the right words to use in conversations. |
|  |  | LANG4 | 4. Communicating thoughts in a conversation. |
|  |  | LANG5 | 5. Following a story in a book or on TV. |
|  |  | LANG6 | 6. Understanding the point of what other people are trying to say. |
|  |  | LANG7 | 7. Remembering the meaning of common words. |
|  |  | LANG8 | 8. Describing a program he/she has watched on TV. |
|  |  | LANG9 | 9. Understanding spoken directions or instructions. |
|  |  | VISSPAT1 | 1. Following a map to find a new location. |
|  |  | VISSPAT2 | 2. Reading a map and helping with directions when someone else is driving. |
|  |  | VISSPAT3 | 3. Finding one's car in a parking lot. |
|  |  | VISSPAT4 | 4. Finding my way back to a meeting spot in the mall or other location. |
|  |  | VISSPAT6 | 5. Finding his/her way around a familiar neighborhood. |
|  |  | VISSPAT7 | 6. Finding his/her way around a familiar store. |
|  |  | VISSPAT8 | 7. Finding his/her way around a house visited many times. |
|  |  | PLAN1 | 1. Planning a sequence of stops on a shopping trip. |
|  |  | PLAN2 | 2. The ability to anticipate weather changes and plan accordingly (i.e., bring a coat or umbrella) |
|  |  | PLAN3 | 3. Developing a schedule in advance of anticipated events. |
|  |  | PLAN4 | 4. Thinking things through before acting. |
|  |  | PLAN5 | 5. Thinking ahead. |
|  |  | ORGAN1 | 1. Keeping living and work space organized. |
|  |  | ORGAN2 | 2. Balancing the checkbook without error. |
|  |  | ORGAN3 | 3. Keeping financial records organized. |
|  |  | ORGAN4 | 4. Prioritizing tasks by importance. |
|  |  | ORGAN5 | 5. Keeping mail and papers organized. |
|  |  | ORGAN6 | 6. Using an organized strategy to manage a medication schedule involving multiple medications. |
|  |  | DIVATT1 | 1. The ability to do two things at once. |
|  |  | DIVATT2 | 2. Returning to a task after being interrupted. |
|  |  | DIVATT3 | 3. The ability to concentrate on a task without being distracted by external things in the environment. |
|  |  | DIVATT4 | 4. Cooking or working and talking at the same time. |
| Everyday cognition - participant |  |  |  |
| ECOGPT - (838, 39) - ['bl', 'm06', 'm12'] |  | MEMORY1 | 1. Remembering a few shopping items without a list. |
|  |  | MEMORY2 | 2. Remembering things that happened recently (such as recent outings, events in the news). |
|  |  | MEMORY3 | 3. Recalling conversations a few days later. |
|  |  | MEMORY4 | 4. Remembering where I have placed objects. |
|  |  | MEMORY5 | 5. Repeating stories and/or questions. |
|  |  | MEMORY6 | 6. Remembering the current date or day of the week. |
|  |  | MEMORY7 | 7. Remembering I have already told someone something. |
|  |  | MEMORY8 | 8. Remembering appointments, meetings, or engagements. |
|  |  | LANG1 | 1. Forgetting the names of objects. |
|  |  | LANG2 | 2. Verbally giving instructions to others. |
|  |  | LANG3 | 3. Finding the right words to use in conversations. |
|  |  | LANG4 | 4. Communicating thoughts in a conversation. |
|  |  | LANG5 | 5. Following a story in a book or on TV. |
|  |  | LANG6 | 6. Understanding the point of what other people are trying to say. |
|  |  | LANG7 | 7. Remembering the meaning of common words. |
|  |  | LANG8 | 8. Describing a program I have watched on TV. |
|  |  | LANG9 | 9. Understanding spoken directions or instructions. |
|  |  | VISSPAT1 | 1. Following a map to find a new location. |
|  |  | VISSPAT2 | 2. Reading a map and helping with directions when someone else is driving. |
|  |  | VISSPAT3 | 3. Finding my car in a parking lot. |
|  |  | VISSPAT4 | 4. Finding my way back to a meeting spot in the mall or other location. |
|  |  | VISSPAT6 | 5. Finding my way around a familiar neighborhood. |
|  |  | VISSPAT7 | 6. Finding my way around a familiar store. |
|  |  | VISSPAT8 | 7. Finding my way around a house visited many times. |
|  |  | PLAN1 | 1. Planning a sequence of stops on a shopping trip. |
|  |  | PLAN2 | 2. The ability to anticipate weather changes and plan accordingly (i.e., bring a coat or umbrella) |
|  |  | PLAN3 | 3. Developing a schedule in advance of anticipated events. |
|  |  | PLAN4 | 4. Thinking things through before acting. |
|  |  | PLAN5 | 5. Thinking ahead. |
|  |  | ORGAN1 | 1. Keeping living and work space organized. |
|  |  | ORGAN2 | 2. Balancing the checkbook without error. |
|  |  | ORGAN3 | 3. Keeping financial records organized. |
|  |  | ORGAN4 | 4. Prioritizing tasks by importance. |
|  |  | ORGAN5 | 5. Keeping mail and papers organized. |
|  |  | ORGAN6 | 6. Using an organized strategy to manage a medication schedule involving multiple medications. |
|  |  | DIVATT1 | 1. The ability to do two things at once. |
|  |  | DIVATT2 | 2. Returning to a task after being interrupted. |
|  |  | DIVATT3 | 3. The ability to concentrate on a task without being distracted by external things in the environment. |
|  |  | DIVATT4 | 4. Cooking or working and talking at the same time. |
| Functional Assessment Questionnaire |  |  |  |
| FAQTOTAL - (1637, 1) - ['bl', 'm06', 'm12'] |  | FAQFINAN | 1. Writing checks, paying bills, or balancing checkbook. |
|  |  | FAQFORM | 2. Assembling tax records, business affairs, or other papers. |
|  |  | FAQSHOP | 3. Shopping alone for clothes, household necessities, or groceries. |
|  |  | FAQGAME | 4. Playing a game of skill such as bridge or chess, working on a hobby. |
|  |  | FAQBEVG | 5. Heating water, making a cup of coffee, turing off the stove. |
|  |  | FAQMEAL | 6. Preparing a balanced meal. |
|  |  | FAQEVENT | 7. Keeping track of current events. |
|  |  | FAQTV | 8. Paying attention to and understanding a TV program, book, or magazine. |
|  |  | FAQREM | 9. Remembering appointments, family occasions, holidays, medications. |
|  |  | FAQTRAVL | 10. Traveling out of the neighborhood, driving, or arranging to take public transportation. |
|  | We only use the total score | FAQTOTAL | Total Score |

**S3 Figure:** **Number of observations available for each feature before and after data imputation**

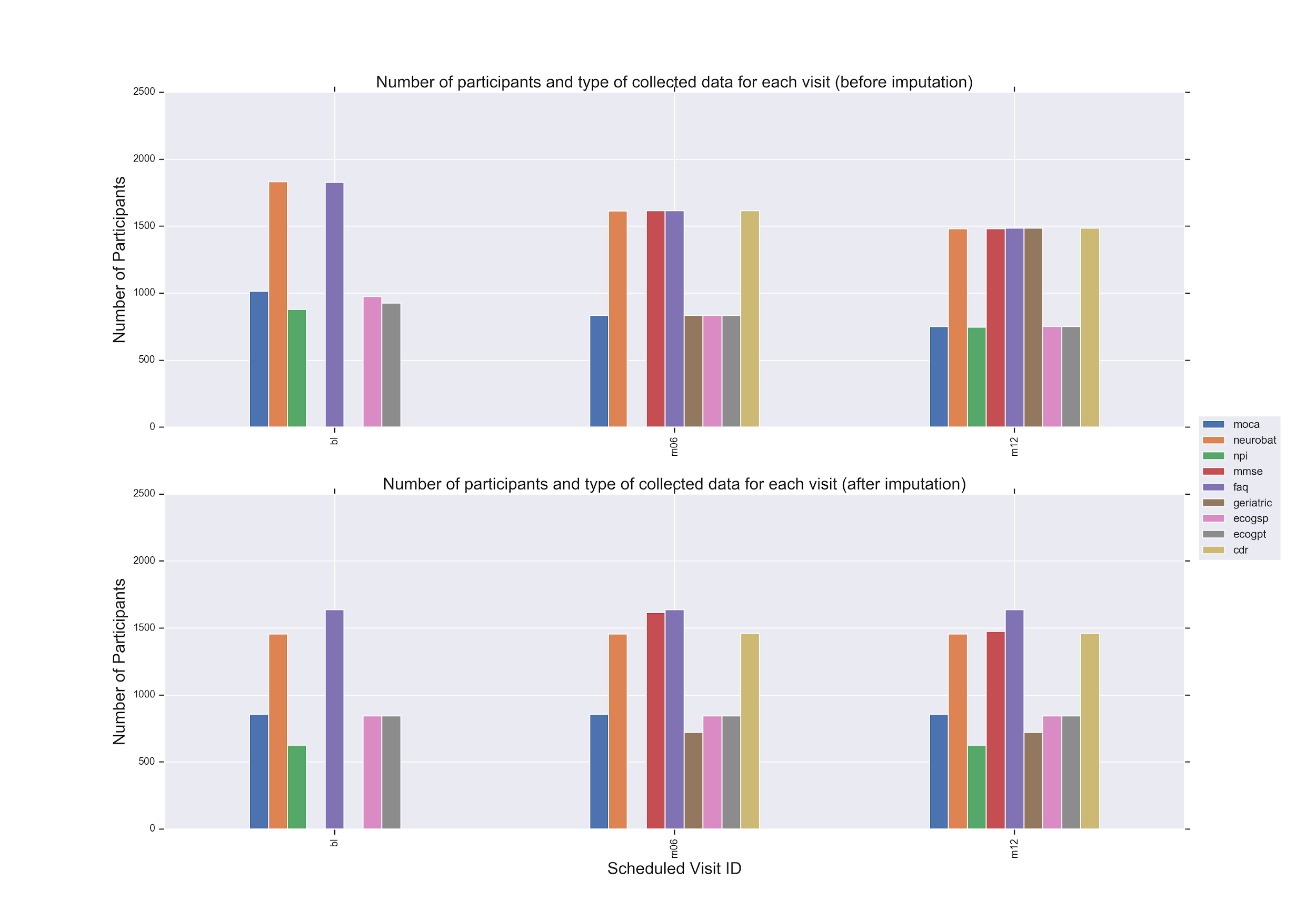

**S4 Table: Axis labels for features using 2D non-negative matrix factorization and Gaussian mixture model**

| **Feature Name** | **Axis Label** | **Description** | **Dataset** |
| --- | --- | --- | --- |
| CATANINTR | 2 | Category Fluency (Animals) - Intrusions | NEUROBAT - m12 |
| TRAAERROM | 2 | Errors of Omission | NEUROBAT - bl |
| NPIFTOT | 2 | Elation/Euphoria: Item score | NPIALL - m12 |
| TRAAERROM | 2 | Errors of Omission | NEUROBAT - m06 |
| NPIBTOT | 2 | Hallucinations: Item score | NPIALL - m12 |
| NPIBTOT | 2 | Hallucinations: Item score | NPIALL - bl |
| NPIFTOT | 2 | Elation/Euphoria: Item score | NPIALL - bl |
| TRAAERROM | 2 | Errors of Omission | NEUROBAT - m12 |
| CATANINTR | 2 | Category Fluency (Animals) - Intrusions | NEUROBAT - bl |
| NPIATOT | 2 | Delusions: Item score | NPIALL - bl |
| CATANINTR | 2 | Category Fluency (Animals) - Intrusions | NEUROBAT - m06 |
| TRAAERRCOM | 2 | Errors of Omission | NEUROBAT - m12 |
| NPIATOT | 2 | Delusions: Item score | NPIALL - m12 |
| NPIHTOT | 2 | Disinhibition: Item score | NPIALL - m12 |
| NPIHTOT | 2 | Disinhibition: Item score | NPIALL - bl |
| AVERR3 | 2 | Total Intrusions (Drum, Curtain, Bell, Coffee, etc.), Trial 3 | NEUROBAT - bl |
| NPIJTOT | 2 | Aberrant Motor Behavior: Item score | NPIALL - bl |
| BNTCSTIM | 2 | Total correct with a semantic cue | NEUROBAT - m12 |
| TRAAERRCOM | 2 | Errors of Omission | NEUROBAT - m06 |
| NPIJTOT | 2 | Aberrant Motor Behavior: Item score | NPIALL - m12 |
| BNTCSTIM | 2 | Total correct with a semantic cue | NEUROBAT - m06 |
| AVERR1 | 2 | Total Intrusions (Drum, Curtain, Bell, Coffee, etc.), Trial 1 | NEUROBAT - m12 |
| AVERR5 | 2 | Total Intrusions (Drum, Curtain, Bell, Coffee, etc.), Trial 5 | NEUROBAT - m12 |
| AVERR3 | 2 | Total Intrusions (Drum, Curtain, Bell, Coffee, etc.), Trial 3 | NEUROBAT - m12 |
| AVERR4 | 2 | Total Intrusions (Drum, Curtain, Bell, Coffee, etc.), Trial 4 | NEUROBAT - m12 |
| NPILTOT | 2 | Appetite and eating disorders: Item score | NPIALL - m12 |
| AVERR4 | 2 | Total Intrusions (Drum, Curtain, Bell, Coffee, etc.), Trial 1 | NEUROBAT - m06 |
| AVERR2 | 2 | Total Intrusions (Drum, Curtain, Bell, Coffee, etc.), Trial 2 | NEUROBAT - m12 |
| TRABERROM | 2 | Errors of Omission | NEUROBAT - m12 |
| NPIKTOT | 2 | Sleep: Item score | NPIALL - m12 |
| BNTCSTIM | 2 | Total correct with a semantic cue | NEUROBAT - bl |
| TRAAERRCOM | 2 | Errors of Commission | NEUROBAT - bl |
| AVERR2 | 2 | Total Intrusions (Drum, Curtain, Bell, Coffee, etc.), Trial 2 | NEUROBAT - m06 |
| AVERR4 | 2 | Total Intrusions (Drum, Curtain, Bell, Coffee, etc.), Trial 4 | NEUROBAT - bl |
| AVERR6 | 2 | Total Intrusions (Drum, Curtain, Bell, Coffee, etc.), Trial 6 | NEUROBAT - m12 |
| NPIKTOT | 2 | Sleep: Item score | NPIALL - bl |
| NPIETOT | 2 | Anxiety: Item score | NPIALL - m12 |
| AVERR5 | 2 | Total Intrusions (Drum, Curtain, Bell, Coffee, etc.), Trial 5 | NEUROBAT - bl |
| AVERR1 | 2 | Total Intrusions (Drum, Curtain, Bell, Coffee, etc.), Trial 1 | NEUROBAT - bl |
| NPICTOT | 2 | Agitation/Aggression: Item score | NEUROBAT -m06 |
| NPIETOT | 2 | Anxiety: Item score | NPIALL - bl |
| TRABERROM | 2 | Errors of Omission | NEUROBAT -m06 |
| AVERR2 | 2 | Total Intrusions (Drum, Curtain, Bell, Coffee, etc.), Trial 2 | NEUROBAT -bl |
| AVERR5 | 2 | Total Intrusions (Drum, Curtain, Bell, Coffee, etc.), Trial 5 | NEUROBAT -m06 |
| NPIDTOT | 2 | Depression/Dysphoria: Item score | NPIALL - bl |
| AVERR1 | 2 | Total Intrusions (Drum, Curtain, Bell, Coffee, etc.), Trial 1 | NEUROBAT -m06 |
| AVERR3 | 2 | Total Intrusions (Drum, Curtain, Bell, Coffee, etc.), Trial 3 | NEUROBAT -m06 |
| NPICTOT | 2 | Agitation/Aggression: Item score | NPIALL - m12 |
| NPIITOT | 2 | Irritability/Lability: Item score | NPIALL - m12 |
| NPILTOT | 2 | Appetite and eating disorders: Item score | NPIALL - m12 |
| AVERR6 | 2 | Total Intrusions | NEUROBAT -m06 |
| NPIITOT | 2 | Irritability/Lability: Item score | NPIALL - bl |
| BNTCPHON | 2 | Number of correct responses following a phonemic cue | NEUROBAT - bl |
| BNTCPHON | 2 | Number of correct responses following a phonemic cue | NEUROBAT -m06 |
| AVERR6 | 2 | Total Intrusions (Drum, Curtain, Bell, Coffee, etc.), Trial 6 | NEUROBAT - bl |
| BNTCPHON | 2 | Number of correct responses following a phonemic cue | NEUROBAT -m12 |
| CATANPERS | 2 | Category Fluency (Animals) - Perseverations | NEUROBAT -m06 |
| TRABERRCOM | 2 | Errors of Commission | NEUROBAT -m12 |
| TRABERROM | 2 | Errors of Omission | NEUROBAT -m12 |
| NPIDTOT | 2 | Depression/Dysphoria: Item score | NPIALL - m12 |
| AVERRB | 2 | Total Intrusions | NEUROBAT - bl |
| NPIGTOT | 2 | Apathy/Indifference: Item score | NPIALL - m12 |
| NPIGTOT | 2 | Apathy/Indifference: Item score | NPIALL - bl |
| CATANPERS | 2 | Category Fluency (Animals) - Perseverations | NEUROBAT - bl |
| BNTSTIM | 2 | Total semantic cues given | NEUROBAT - m12 |
| GDTOTAL | 2 | Total Depression Score | GDSCALE - m06 |
| GDTOTAL | 2 | Total Depression Score | GDSCALE - m12 |
| BNTSTIM | 2 | Total semantic cues given | NEUROBAT - m06 |
| CATANPERS | 2 | Category Fluency (Animals) - Perseverations | NEUROBAT - m12 |
| BNTSTIM | 2 | Total semantic cues given | NEUROBAT - bl |
| AVERRB | 2 | Total Intrusions | NEUROBAT - m06 |
| TRABERRCOM | 2 | Errors of Commission | NEUROBAT - bl |
| AVDELERR2 | 2 | Total Intrusions (Drum, Curtain, Bell, Coffee, etc.), Trial 2 | NEUROBAT - bl |
| AVERRB | 2 | Total Intrusions | NEUROBAT - m12 |
| vis | 2 | Following a map to find a new location, Reading a map and helping with directions when someone else is driving., Finding one's car in a parking lot.  , Finding my way back to a meeting spot in the mall or other location., etc. | ECOGPT - m06 |
| vis | 2 | Following a map to find a new location, Reading a map and helping with directions when someone else is driving., Finding one's car in a parking lot.  , Finding my way back to a meeting spot in the mall or other location., etc. | ECOGPT - bl |
| TRABERRCOM | 2 | Errors of Commission | NEUROBAT - m06 |
| AVDELERR2 | 2 | Total Intrusions (Drum, Curtain, Bell, Coffee, etc.), Trial 2 | NEUROBAT - m06 |
| plan | 2 | Planning a sequence of stops on a shopping trip, The ability to anticipate weather changes and plan accordingly (i.e., bring a coat or umbrella), Developing a schedule in advance of anticipated events, etc. | ECOGPT - bl |
| AVDELERR2 | 2 | Total Intrusions (Drum, Curtain, Bell, Coffee, etc.), Trial 2 | NEUROBAT - m12 |
| org | 2 | Keeping living and work space organized, Balancing the checkbook without error.  , Keeping financial records organized., Prioritizing tasks by importance.  , etc. | ECOGPT - bl |
| BNTPHON | 2 | Total phonemic cues given | NEUROBAT - bl |
| org | 2 | Keeping living and work space organized, Balancing the checkbook without error.  , Keeping financial records organized., Prioritizing tasks by importance.  , etc. | ECOGPT - m06 |
| TRAASCOR | 2 | Part B - Time to complete | NEUROBAT - bl |
| BNTPHON | 2 | Total phonemic cues given | NEUROBAT - m06 |
| TRAASCOR | 2 | Part B - Time to complete | NEUROBAT - m06 |
| BNTPHON | 2 | Total phonemic cues given | NEUROBAT - m12 |
| FAQ | 2 | FAQ Total Score | FAQ - bl |
| lang | 2 | Forgetting the names of objects, Verbally giving instructions to others, Finding the right words to use in conversations, Communicating thoughts in a conversation, Following a story in a book or on TV. etc. | ECOGPT - m06 |
| lang | 2 | Forgetting the names of objects, Verbally giving instructions to others, Finding the right words to use in conversations, Communicating thoughts in a conversation, Following a story in a book or on TV. etc. | ECOGPT - bl |
| TRAASCOR | 2 | Part B - Time to complete | NEUROBAT - m12 |
| division | 2 | The ability to do two things at once, Returning to a task after being interrupted, the ability to concentrate on a task without being distracted by external things in the environment, etc. | ECOGPT - bl |
| CDCARE | 2 | Personal Care Score | CDR - m06 |
| FAQ | 2 | Total Score | FAQ - m06 |
| CDHOME | 2 | Home and Hobbies Score | CDR - m06 |
| CDCARE | 2 | Personal Care Score | CDR - m12 |
| FAQ | 2 | FAQ Total Score | FAQ - m12 |
| CDHOME | 2 | Home and Hobbies Score | CDR - m12 |
| memory | 2 | Remembering a few shopping items without a list. | ECOGPT - m12 |
| memory | 2 | Remembering a few shopping items without a list. | ECOGPT - m06 |
| memory | 2 | Remembering a few shopping items without a list. | ECOGPT - bl |
| CDCOMMUN | 2 | Community Affairs Score | CDR - m06 |
| CDORIENT | 2 | Orientation Score | CDR - m06 |
| CDJUDGE | 2 | Judgment and Problem Solving Score | CDR - m06 |
| CDCOMMUN | 2 | Community Affairs Score | CDR - m12 |
| TRABSCOR | 2 | Part A - Time to Complete | NEUROBAT - bl |
| CDORIENT | 2 | Orientation Score | CDR - m12 |
| CDJUDGE | 2 | Judgment and Problem Solving Score | CDR - m12 |
| CDMEMORY | 2 | Memory Score | CDR - m06 |
| TRABSCOR | 2 | Part A - Time to Complete | NEUROBAT - m06 |
| CDMEMORY | 2 | Memory Score | CDR - m12 |
| TRABSCOR | 2 | Part A - Time to Complete | NEUROBAT - m12 |
| AVDEL30MIN | 1 | 30 Minute Delay Total | NEUROBAT - bl |
| AVDEL30MIN | 1 | 30 Minute Delay Total | NEUROBAT - m12 |
| AVDEL30MIN | 1 | 30 Minute Delay Total | NEUROBAT - m06 |
| LDELTOTAL | 1 | Total Number of Story Units Recalled | NEUROBAT - m12 |
| AVTOT6 | 1 | Total Intrusions (Drum, Curtain, Bell, Coffee, etc.) Trial 6 Total | NEUROBAT - m06 |
| AVTOT6 | 1 | Total Intrusions (Drum, Curtain, Bell, Coffee, etc.) Trial 6 Total | NEUROBAT - m06 |
| AVTOT6 | 1 | Total Intrusions (Drum, Curtain, Bell, Coffee, etc.) Trial 6 Total | NEUROBAT - bl |
| AVDELERR1 | 1 | Total Intrusions (Drum, Curtain, Bell, Coffee, etc.) | NEUROBAT - m12 |
| LIMMTOTAL | 1 | Total Number of Story Units Recalled | NEUROBAT - m12 |
| AVTOT3 | 1 | Total Intrusions (Drum, Curtain, Bell, Coffee, etc.) Trial 3 Total | NEUROBAT - bl |
| AVTOT5 | 1 | Total Intrusions (Drum, Curtain, Bell, Coffee, etc.) Trial 5 Total | NEUROBAT - m12 |
| AVDELERR1 | 1 | Total Intrusions (Drum, Curtain, Bell, Coffee, etc.), Trial 1 | NEUROBAT - m06 |
| AVTOT4 | 1 | Total Intrusions (Drum, Curtain, Bell, Coffee, etc.)Trial 4 Total | NEUROBAT - bl |
| AVTOTB | 1 | List B Total | NEUROBAT - m12 |
| AVTOT5 | 1 | Total Intrusions (Drum, Curtain, Bell, Coffee, etc.) Trial 5 Total | NEUROBAT - bl |
| AVTOT4 | 1 | Total Intrusions (Drum, Curtain, Bell, Coffee, etc.) Trial 4 Total | NEUROBAT - m06 |
| AVDELERR1 | 1 | Total Intrusions (Drum, Curtain, Bell, Coffee, etc.), Trial 1 | NEUROBAT - bl |
| AVTOTB | 1 | Total Intrusions (Drum, Curtain, Bell, Coffee, etc.) List B Total | NEUROBAT - bl |
| AVTOT1 | 1 | Trial 1 Total (Time to complete) | NEUROBAT - m12 |
| AVTOT3 | 1 | Total Intrusions (Drum, Curtain, Bell, Coffee, etc.)Trial 3 Total | NEUROBAT - m12 |
| AVTOT2 | 1 | Total Intrusions (Drum, Curtain, Bell, Coffee, etc.) Trial 2 Total | NEUROBAT - m12 |
| AVTOT4 | 1 | Total Intrusions (Drum, Curtain, Bell, Coffee, etc.) Trial 4 Total | NEUROBAT - m12 |
| AVTOT5 | 1 | Total Intrusions (Drum, Curtain, Bell, Coffee, etc.) Trial 5 Total | NEUROBAT - m06 |
| AVTOT1 | 1 | Total Intrusions (Drum, Curtain, Bell, Coffee, etc.) Trial 1 Total | NEUROBAT - m06 |
| AVTOT3 | 1 | Total Intrusions (Drum, Curtain, Bell, Coffee, etc.) Trial 3 Total | NEUROBAT - m06 |
| AVTOTB | 1 | Total Intrusions (Drum, Curtain, Bell, Coffee, etc.) List B Total | NEUROBAT - m06 |
| AVTOT2 | 1 | Total Intrusions (Drum, Curtain, Bell, Coffee, etc.) Trial 2 Total | NEUROBAT - m06 |
| AVTOT2 | 1 | Total Intrusions (Drum, Curtain, Bell, Coffee, etc.) Trial 2 Total | NEUROBAT - bl |
| CATANIMSC | 1 | Category Fluency (Animals) - Total Correct | NEUROBAT - bl |
| AVTOT1 | 1 | Total Intrusions (Drum, Curtain, Bell, Coffee, etc.) Trial 1 Total | NEUROBAT - m12 |
| CATANIMSC | 1 | Category Fluency (Animals) - Total Correct | NEUROBAT - m12 |
| CATANIMSC | 1 | Category Fluency (Animals) - Total Correct | NEUROBAT - m06 |
| attention | 1 | Attention to digits and letters | MOCA - m06 |
| attention | 1 | Attention to digits and letters | MOCA - bl |
| AVDELTOT | 1 | Recognition Score | NEUROBAT - m12 |
| fluency | 1 | Letter Fluency - F: Total number of correct words | MOCA - m12 |
| AVDELTOT | 1 | Recognition Score | NEUROBAT - m06 |
| fluency | 1 | Letter Fluency - F: Total number of correct words | MOCA - m06 |
| fluency | 1 | Letter Fluency - F: Total number of correct words | MOCA - bl |
| AVDELTOT | 1 | Recognition Score | NEUROBAT - bl |
| sen repetetion | 1 | Sentence Repetetion | MOCA - m12 |
| visuosoconstructional | 1 | Visioconsytructional (Draw Cube , clock etc) | MOCA - m12 |
| sen repetetion | 1 | Sentence Repetetion | MOCA - m06 |
| delayed word recall | 1 | Delayed word recall | MOCA - m12 |
| trail making | 1 | Alternating Trail Making | MOCA - bl |
| delayed word recall | 1 | Delayed word recall | MOCA - bl |
| trail making | 1 | Alternating Trail Making | MOCA - m12 |
| delayed word recall | 1 | Delayed word recall | MOCA - m06 |
| visuosoconstructional | 1 | Visioconsytructional (Draw Cube , clock etc) | MOCA - m06 |
| mmse MMSCORE | 1 | MMSE TOTAL SCORE | MMSE - m06 |
| CLOCKSCOR | 1 | Total clock score | MMSE - m06 |
| sen repetetion | 1 | Sentence Repetetion | MOCA - bl |
| mmse MMSCORE | 1 | MMSE TOTAL SCORE | MOCA - m12 |
| abstraction | 1 | Abstraction (train-bicycle, watch-ruler) | MOCA - m12 |
| BNTSPONT | 1 | Total correct without a cue | NEUROBAT - bl |
| abstraction | 1 | Abstraction (train-bicycle, watch-ruler) | MOCA - bl |
| trail making | 1 | Alternating Trail Making | MOCA - m06 |
| BNTSPONT | 1 | Total correct without a cue | NEUROBAT - m06 |
| CLOCKSCOR | 1 | Total clock score | NEUROBAT - bl |
| BNTSPONT | 1 | Total correct without a cue | NEUROBAT - m12 |
| visuosoconstructional | 1 | Visioconsytructional (Draw Cube , clock etc) | NEUROBAT - m12 |
| abstraction | 1 | Abstraction (train-bicycle, watch-ruler) | MOCA - m06 |
| orientation | 1 | Orientation (Date,Month, Year,Day,Place,city) | MOCA - m12 |
| immediate recall | 1 | Immediate word recall | MOCA - m06 |
| orientation | 1 | Orientation (Date,Month, Year,Day,Place,city) | MOCA - bl |
| orientation | 1 | Orientation (Date,Month, Year,Day,Place,city) | MOCA - m06 |
| CLOCKSCOR | 1 | Total clock score | NEUROBAT - m06 |
| immediate recall | 1 | Immediate word recall | MOCA - m12 |
| immediate recall | 1 | Immediate word recall | MOCA - bl |
| naming | 1 | Naming (Lion, Rhino, camel) | MOCA - m12 |
| COPYSCOR | 1 | Total copy score | NEUROBAT - m12 |
| moca naming | 1 | Naming (Lion, Rhino, camel) | MOCA - m06 |
| moca naming | 1 | Naming (Lion, Rhino, camel) | MOCA - m06 |
| COPYSCOR | 1 | Total copying score | NEUROBAT - bl |
| COPYSCOR | 1 | Total copy score | NEUROBAT - m06 |

**S5 Figure: Projection mapping for clinical features to new 2-dimensional AD progression space axis. Here, first axis corresponds to cognition related features and second axis corresponds to memory related features.**

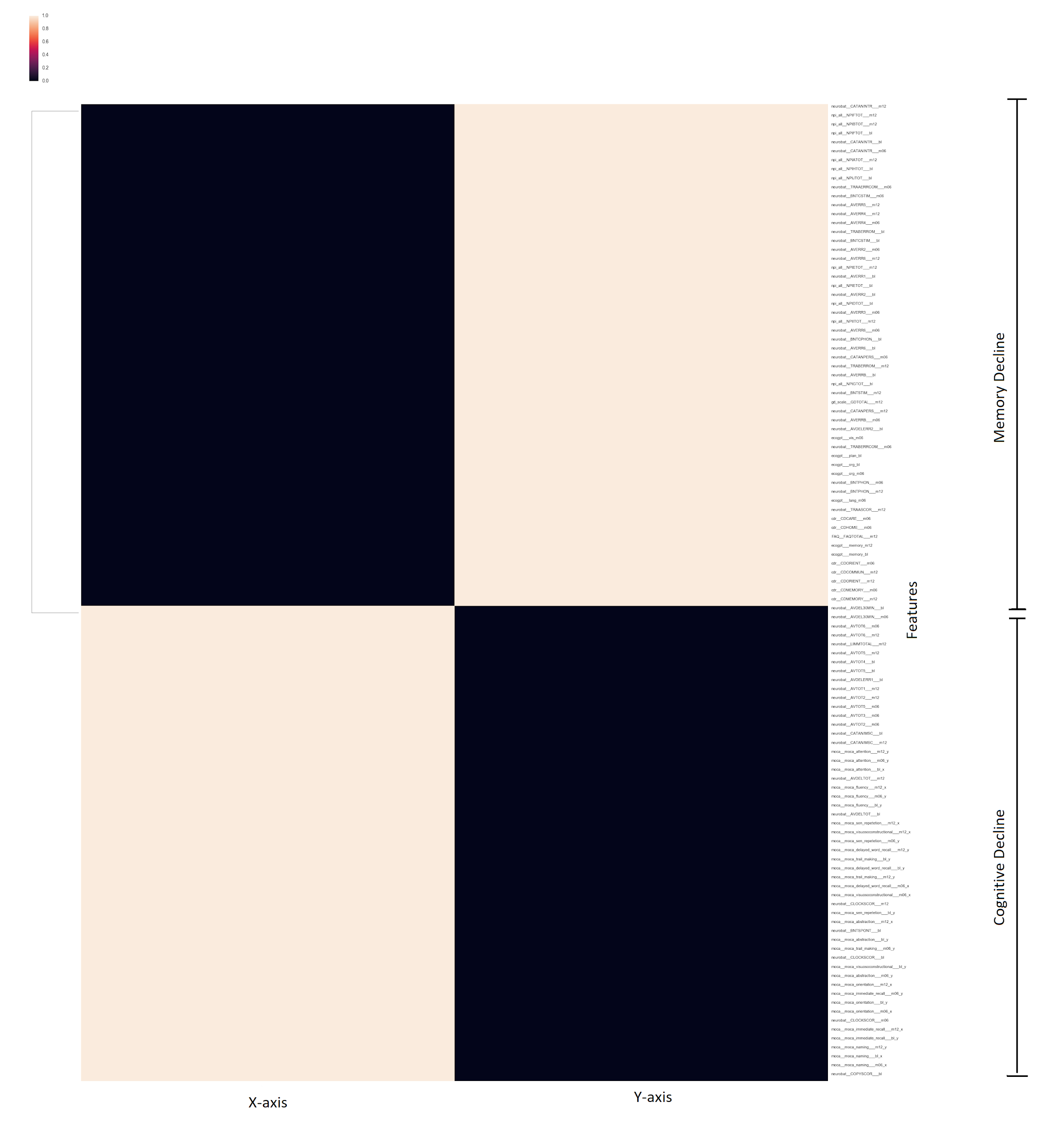

**S6 Figure: Comparison of different algorithms for the prediction of progression after 24 months from the baseline. Five-fold cross-validation accuracy was used to evaluate each model. Random forest algorithm provides the highest accuracy.**

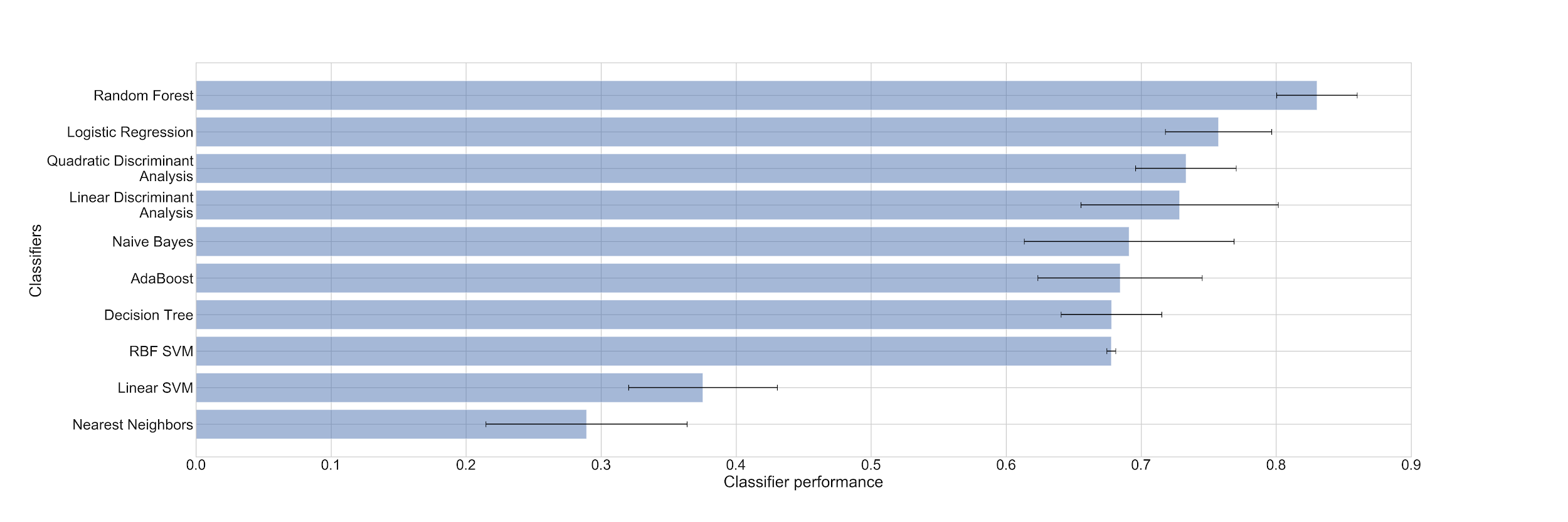

**S7 Figure: Comparison of different algorithms for the prediction of progression after 48 months from the baseline. Five-fold cross-validation accuracy was used to evaluate each model. Random forest algorithm provides the highest accuracy.**

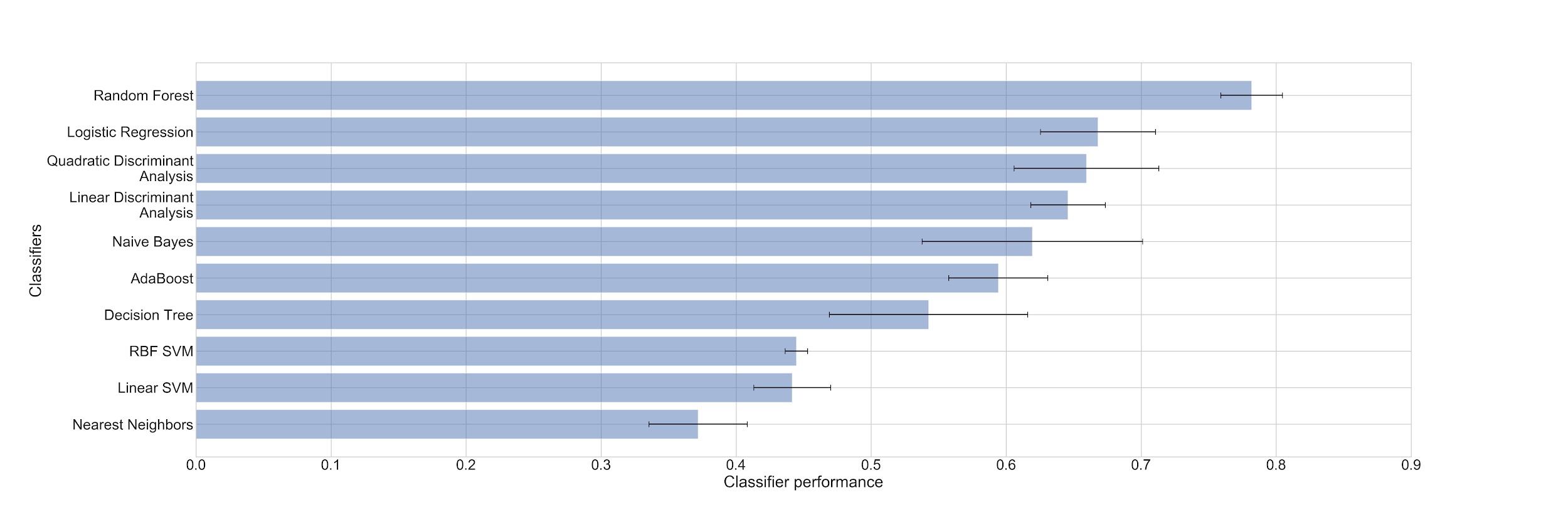

**S8 Figure: Percent share of controls, MCI and dementia patients for different subtypes present after 48 months from baseline. The share of MCI patients is decreasing with increase in progression rate.**

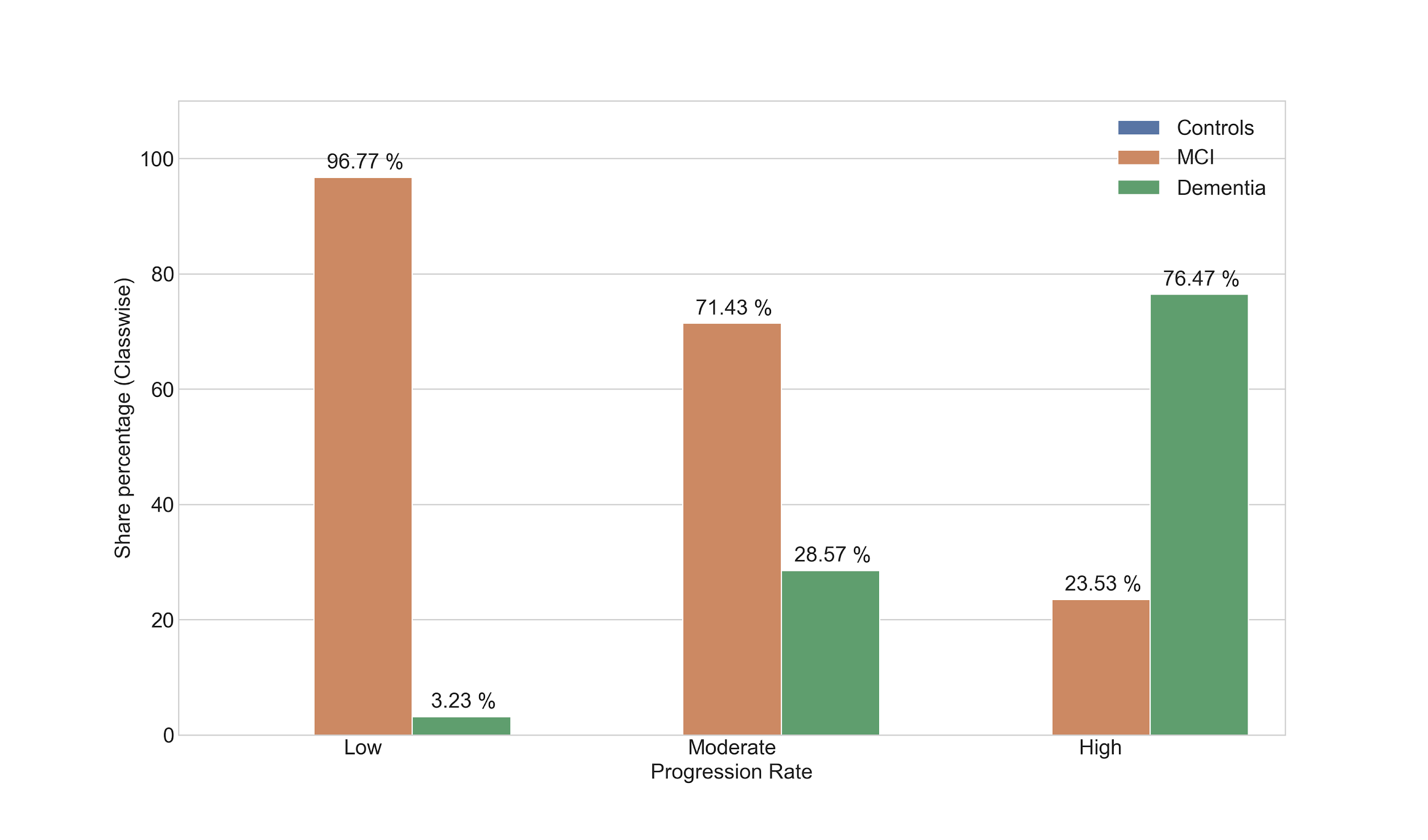

**S9 Figure: Percent share of APOEε4 variants for different subtypes after 48 months from baseline. The share of 0 occurrences of APOEε4 variants is decreasing with increase in progression rate, whereas the share of other two 1 and 2 occurrences is increasing.**

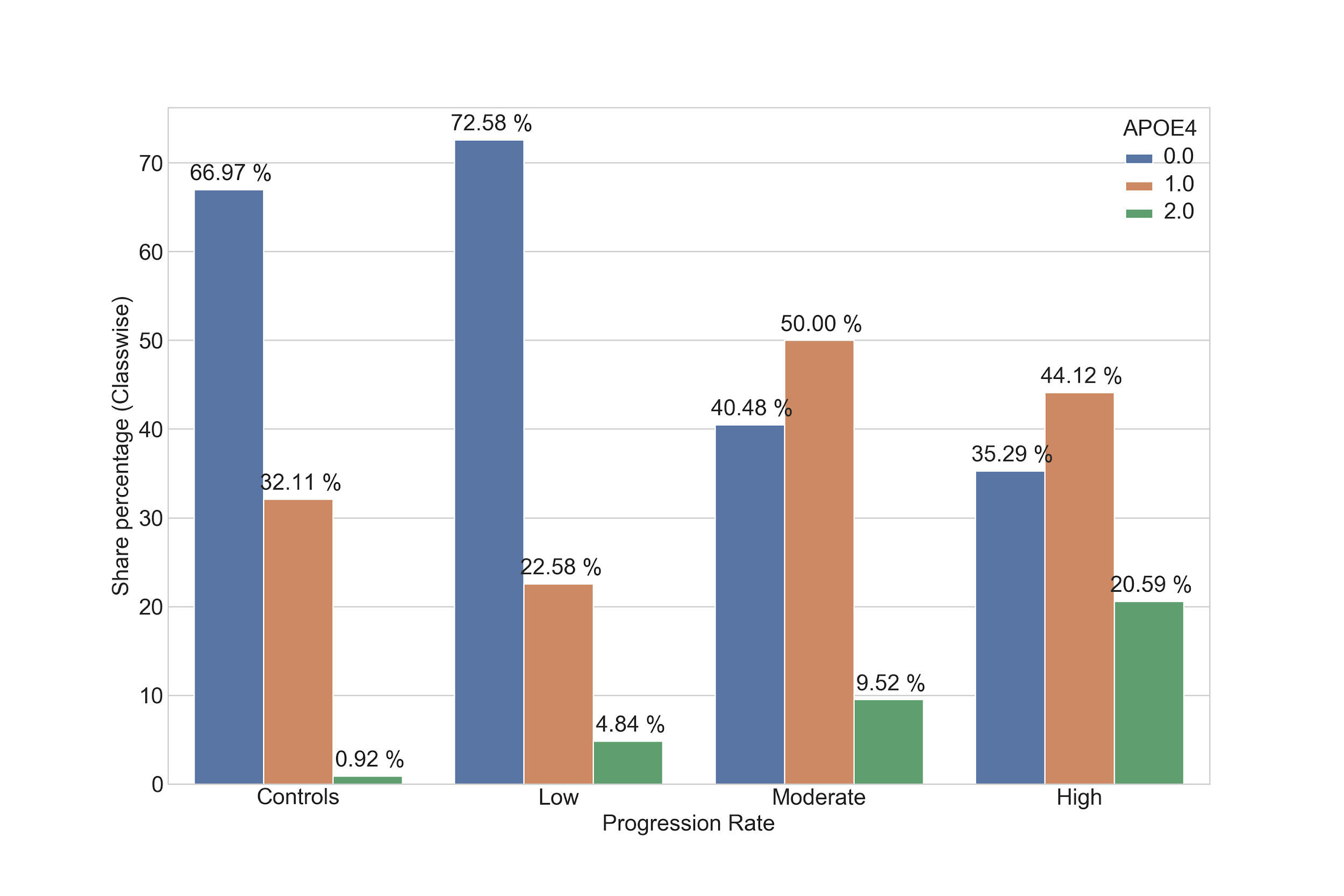

**S10 Figure. The distribution of MMSE score for each AD subtype (subtypes at the 48th month) after 6 and 12 months from the baseline. MMSE score decreases with increase in the progression rate.**

| 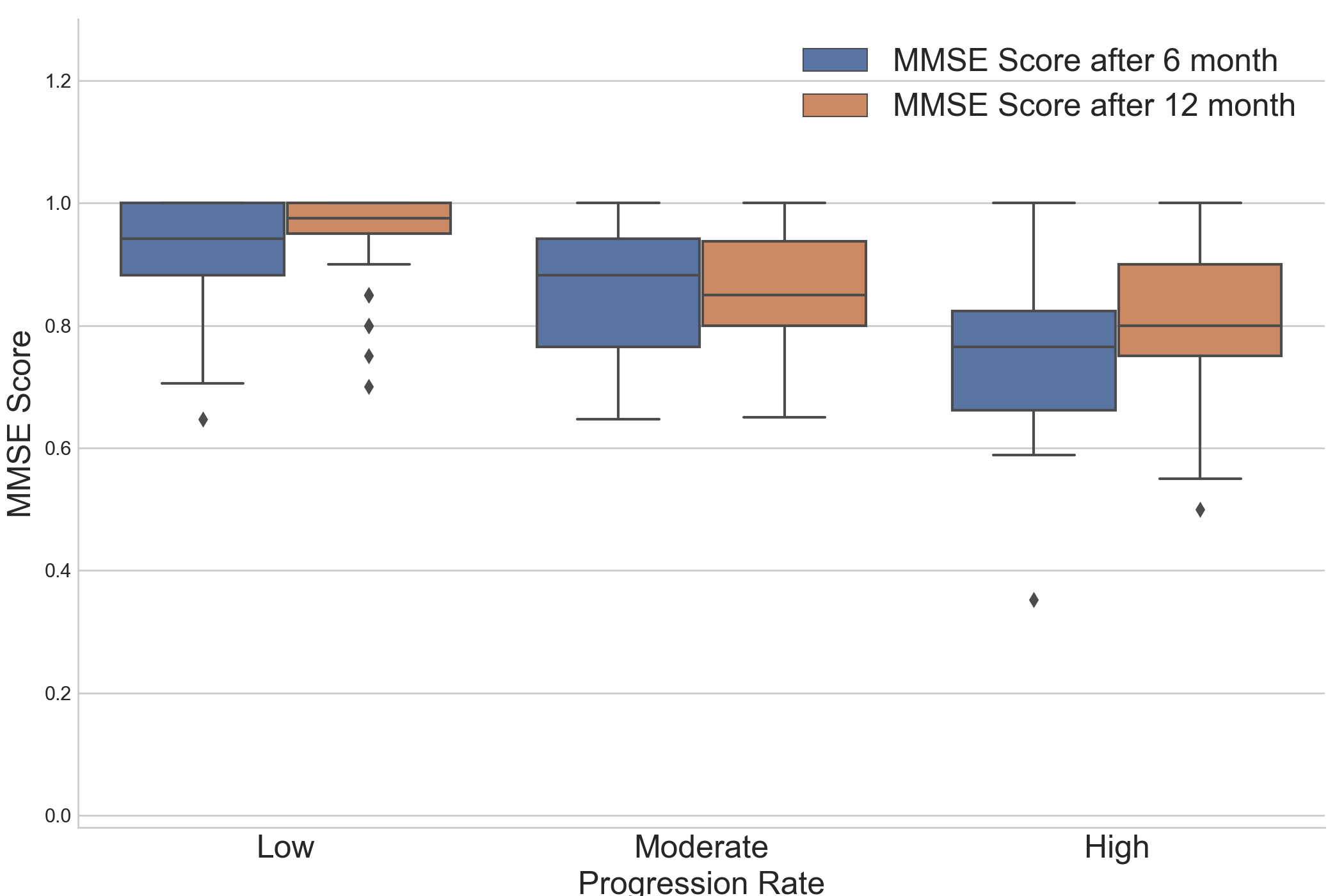  **S11 Figure: The distribution of FAQ total score score for each AD subtype (subtypes at the 48th month) after 6 and 12 months from the baseline. FAQ total score increases with increase in the progression rate.**  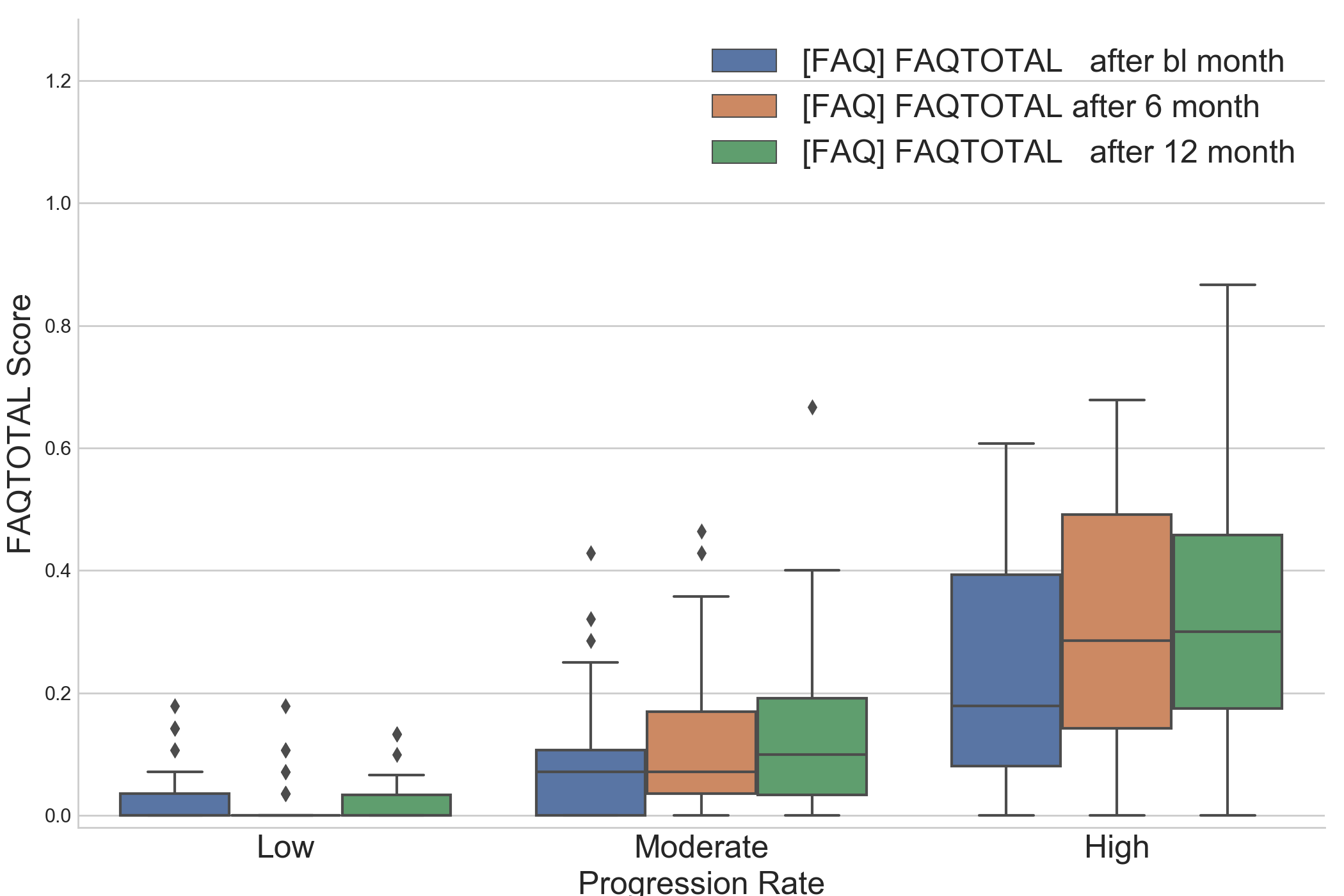 |
| --- |
